# The secreted peptide IRP functions as a DAMP in rice immunity

**DOI:** 10.1101/2021.01.31.429004

**Authors:** Pingyu Wang, Huimin Jia, Ting Guo, Yuanyuan Zhang, Zhengguo Li, Yoji Kawano

## Abstract

Small signaling peptides play important roles in various plant processes, but information regarding their involvement in plant immunity is limited. We previously identified a novel small secreted protein in rice, named immune response peptide (IRP) by the integrated multi-omics analyses. Here, we studied IRP functions in rice immunity. Rice plants overexpressing *IRP* enhanced resistance to the blast fungus. Application of the IRP peptide to rice suspension cells triggered the expression of *IRP* itself and defense gene *PAL1*. RNA-seq results revealed that 84% of genes upregulated by IRP peptide were also induced by chitin, including 13 *OsWRKY* transcription factors, indicating that IRP and chitin share the similar signaling pathway. Co-treatment with chitin and IRP elevated the expression level of *PAL1* and *OsWRKY*s in an additive manner. When results of *IRP* and *PAL1* expression and MAPK activation by IRP were compared with those by chitin, IRP had a stronger effect on MAPK activation rather than *IRP* and *PAL1* expression. Collectively, our findings indicate that IRP functions as a damage-associated molecular pattern (DAMP) in rice immunity, regulating MAPKs and *Os*WRKYs to amplify chitin signaling, and provide new insights into how PAMPs and DAMPs cooperatively regulate rice immunity.

## INTRODUCTION

Plants live in environments where they are exposed to attacks from various pathogens and insect herbivores. To survive, plants have evolved two types of immune systems: pattern-triggered immunity (PTI) and effector-triggered immunity (ETI) (Dodds and Rathjen, 2010; Wang et al., 2017). In the first tier of the plant immune system, PTI, cell surface-localized pattern recognition receptors (PRRs) perceive microbial molecules called pathogen- or microbe-associated molecular patterns (PAMPs or MAMPs). However, pathogens can suppress PTI by injecting virulence molecules called effectors into the plant cell cytoplasm. During co-evolution, plants have developed a second layer of the innate immune system, ETI, which is governed by intracellular receptor disease resistance proteins (Jones and Dangl, 2006; Cui et al., 2015).

PAMPs such as bacterial flagellin, elongation factor Tu, peptidoglycan (PGN) and fungal chitin are essential for pathogen survival and are highly structurally conserved among a wide variety of pathogens (Newman et al., 2013). PRRs are receptor-like kinases (RLKs) or receptor-like proteins (RLPs) with specific extracellular domains for perceiving PAMPs. After the recognition of PAMPs by PRRs, serial immune events are induced, including calcium influx, extracellular alkalinization, reactive oxygen species (ROS) production, mitogen-activated protein kinase (MAPK) activation and global transcriptional reprogramming. Besides, endogenous molecules called damage-associated molecular patterns (DAMPs) are produced to regulate the innate immune system. DAMPs are released from plants upon wounding, microbial infection and herbivory (Hou et al., 2019). Some oligomeric fragments from the plant cell wall such as oligogalacturonides and cutin monomers are identified as DAMPs (Benedetti et al., 2015; Ziv et al., 2018). It has been reported that several endogenous peptides that are processed from longer precursors and secreted into the apoplast also function as DAMPs. Interestingly, PAMPs and DAMPs share similar signaling components (Boller and Felix, 2009). Moreover, DAMPs are thought to function as amplifiers in PTI responses (Yamaguchi et al., 2010; Albert, 2013; Hou et al., 2014; Stegmann et al., 2017).

Increasing numbers of signal peptide families have been identified recently, opening a new perspective in plant immunity. Systemin, the first identified peptide hormone in plants, is an 18-amino acid (aa) peptide produced from its precursor, prosystemin (Pearce et al., 1991; McGurl et al., 1992). It can enhance the synthesis of defense-related proteinase inhibitors and can control jasmonic acid (JA) signaling to regulate the systemic defense response (Li et al., 2003; Li et al., 2006). *Arabidopsis thaliana* Pep1 (*At*Pep1) induces the expression of defense genes and the production of H_2_O_2_. The *AtPep1* precursor gene *PROPEP1* is inducible by wounding, methyl JA and ethylene as well as *At*Pep1 peptide. Constitutive overexpression of *PROPEP1* confers resistance against the root pathogen *Pythium irregular*. *At*Peps are recognized by two leucine-rich repeat (LRR) receptor kinase, PEPR1 and PEPR2 (Yamaguchi et al., 2006; Yamaguchi et al., 2010). The pentapeptide phytosulfokines (PSKs) were originally found to play vital roles in plant growth and development. Previous several studies suggested that PSKs are also involved in plant immunity (Loivamaki et al., 2010; Igarashi et al., 2012; Rodiuc et al., 2016). The expression of *AtPSK2* and *PSK receptor 1* (*PSKR1*) is induced by fungal elicitors and fungal pathogens. PSKs appear to have an antagonistic manner in regulating the immune response in *Arabidopsis* to necrotrophic and biotrophic pathogens by enhancing the JA-mediated signaling but reducing the salicylic acid (SA)-mediated signaling (Mosher and Kemmerling, 2013). Rapid alkalinization factor (RALF) was isolated following the observation of rapid alkalinization in the medium of tobacco suspension cells (Pearce et al., 2001). Later, RALFs were found to play both positive and negative roles in plant immunity (Atkinson et al., 2013; Stegmann et al., 2017). A gene family encoding *PIP*s was identified through an in-silico approach (Hou et al., 2014). The expression of *PIP1* and *PIP2* is induced by a variety of pathogens and PAMPs. Overexpression of *PIP1* and *PIP2* increases pathogen resistance in *Arabidopsis*. RLK7 was identified as a receptor for PIP. We are unaware of any reports of peptide DAMPs in rice, and the downstream signaling of DAMPs is poorly understood.

To isolate peptide DAMPs that are involved in rice immunity, we recently developed a pipeline by combining transcriptomics- and proteomics-based screening (Wang et al., 2020). We identified 236 small secreted proteins including members of two known peptide DAMP families, RALF and PSK, as well as a novel gene, *immune response peptide* (*IRP*). The expression of *IRP* is induced by bacterial PGN and fungal chitin. Overexpressing *IRP* in rice suspension cells promotes the expression of the defense gene phenylalanine ammonia-lyase1 (*PAL1*) and triggers the activation of MAPKs. The protein level of IRP increases in suspension cell medium after chitin treatment. In this study, we further clarified the functions of IRP in rice. Rice plants overexpressing *IRP* displayed increased resistance to blast fungus. IRP peptide treatment enhanced the expression of both its precursor *IRP* and *PAL1*, as well as the activity of MAPKs. Thirteen of fourteen WRKY transcription factors induced by IRP peptide were also triggered by chitin treatment. Collectively, our findings indicate that IRP acts as a DAMP to amplify chitin signaling.

## RESULTS

### IRP positively regulates rice immunity

To investigate the function of IRP in rice immunity, we generated *IRP* overexpression and knockout rice plants. We previously reported that T1 *IRP* overexpression plants exhibited a dwarf phenotype (Wang et al., 2020); however, we also found that T3 *IRP* overexpression plants grew like wildtype plants. We performed an infection assay using T3 *IRP* overexpression plants and measured the lesions induced by a virulent rice blast fungus, *Magnaporthe oryzae* (race 007.0). The *IRP* expression level in T3 overexpression plants was confirmed by qPCR (Figure 1a). Compared with wildtype plants, smaller lesions were observed in T3 plants seven days after inoculation, indicating that the overexpression of IRP enhances disease resistance to the virulent rice blast fungus (Figure 1b, c). *IRP* knockout mutants were generated by the CRISPR/Cas9 system. We obtained two independent mutant lines: *irp #1* has a one-base insertion at the 5’-23^rd^ base, resulting in a frameshift, and *irp #2* has a two-base deletion at the 5’-102^nd^ and 103^rd^ bases, resulting in a premature stop codon (Figure S1a). Neither of these *irp* mutants showed an obvious resistance to rice blast fungus (Figure S1b, c).

**Figure 1.**
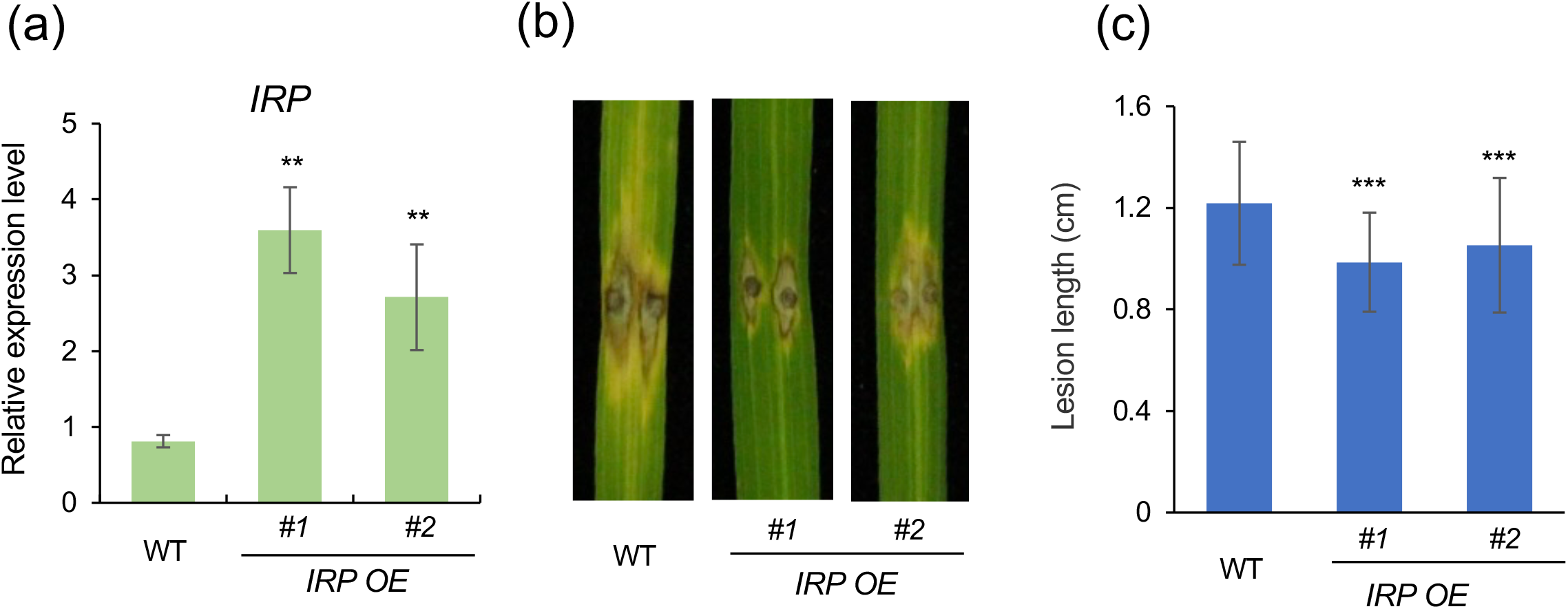
IRP functions in rice immunity. (a) Relative expression level of *IRP* in the overexpression plants was measured by qPCR. Error bars indicate SD (standard deviation). Statistically significant differences between wildtype (WT) and *IRP* overexpression lines (OE) are depicted with asterisks (**, p < 0.01; n = 4) according to the two-tailed *t*-test. (b) Four-week-old rice plants were infected with the fungal pathogen *M. oryzae* and photos of lesions were taken one week after inoculation. (c) Quantitative analysis of lesion size one week after blast fungus infection. Error bars indicate SD. Statistically significant differences between wildtype and overexpression lines are depicted with asterisks (***, p < 0.001, n > 50, two-tailed *t*-test).

### IRP peptide triggers the rice immune response

The *IRP* gene encodes a 66-aa protein containing a 32-aa N-terminal signal sequence (Figure 2a). In general, this signal sequence directs a newly synthesized protein toward the secretory pathway (Matsubayashi, 2014). Previously, we detected a 20-aa peptide corresponding to the C-terminus of IRP in rice suspension cell medium by mass spectrometry analysis, and found that its concentration increases in response to chitin, suggesting that IRP is secreted into the apoplast to regulate rice immunity (Wang et al., 2020). To further analyze its function, we synthesized IRP peptide without the N-terminal signal peptide sequence (Figure 2a). Since the expression of *IRP* and *PAL1* increased to their highest level 1 h after chitin treatment in rice suspension cells, we monitored *IRP* and *PAL1* expression at 1 h after IRP peptide treatment. As a result, IRP peptide was able to induce *IRP* expression at 20 nM. This effect was enhanced as the IRP peptide concentration increased, and induction was maximal at 500 nM (Figure 2b). IRP peptide also induced the expression of *PAL1* in a manner similar to *IRP* (Figure 2c). Time-course analysis revealed that the maximum expression level of *IRP* and *PAL1* occurred at 0.5 h and 1 h after IRP peptide treatment, respectively, and *IRP* expression remained high over a prolonged period (Figure 2d, e). In the previous study, we found that overexpression of *IRP* induces activation of two MAPKs, MPK3 and MPK6, in suspension cells (Wang et al., 2020). Therefore, the MAPK activities after IRP treatment were checked. The results indicated that MPK3/MPK6 were activated 5 min after IRP peptide treatment and their activity peaked at 15 min (Figure 2f). In addition, MPK3/MPK6 activation reached a plateau at 500 nM IRP peptide treatment (Figure 2g). On the other hand, unlike chitin, IRP peptide did not trigger ROS production (Figure 2h, i). The ROS production in suspension cells co-treated with chitin and IRP was not visibly different from chitin treatment alone.

**Figure 2.**
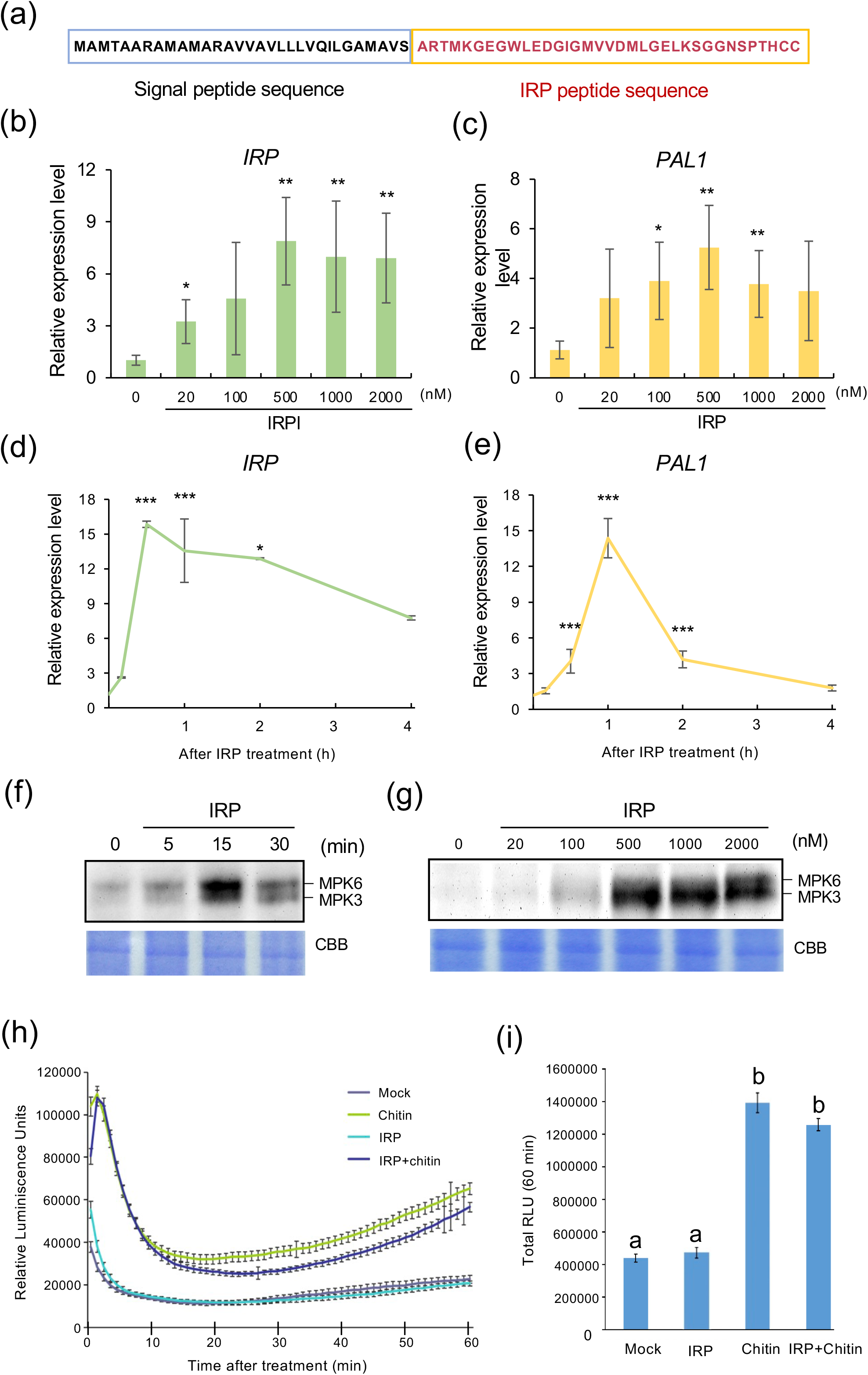
IRP peptide induces immune responses. (a) Amino acid sequence of IRP. Black letters indicate the signal peptide sequence and red letters indicate the C-terminal IRP peptide sequence. (b, c) Relative expression level of *IRP* (b) and *PAL1* (c) in suspension cells treated with different concentrations of IRP peptide for 1 h. Error bars indicate SD. Statistically significant differences between treatment and mock are depicted with asterisks (*p < 0.05; **p < 0.01, n=4, two-tailed *t*-test). (d, e) Relative expression of *IRP* (d) and *PAL1* (e) in rice suspension cells treated with 500 nM IRP peptide for the indicated time period. Error bars indicate SD. Statistically significant differences between treatment and mock are depicted with asterisks (*, p < 0.05; ***, p < 0.001, n=3, two-tailed *t*-test). (f, g) IRP peptide induces MAPK activation. MAPK activation was detected at the indicated times of treatment with 500 nM IRP peptide (f) and at different concentrations of IRP peptide for a 15-min treatment (g). Three independent experiments were performed and similar results were obtained. CBB, Coomassie Brilliant Blue staining. (h, i) ROS production by IRP peptide in rice suspension cells. ROS production at 1-min intervals (h) and total ROS production in 1 h (i) were determined in suspension cells treated with 2 μg/mL chitin, 1 μM IRP or 2 μg/mL chitin and 1 μM IRP. RLU, relative light units. Error bars indicate SD. Statistically significant differences between treatment and mock are depicted with asterisks (*p* < 0.05, n=17, two-tailed *t*-test).

To investigate the immune response induced by IRP, an RNA-seq analysis was performed. In total, 767 differentially expressed genes (DEGs) (log_2_FC > 1 in expression, FDR < 0.05) under IRP treatment were identified, including 695 upregulated and 72 downregulated genes (Table S1). Gene ontology (GO) analysis indicated that IRP-upregulated genes were enriched for molecular function including protein serine/threonine kinase activity and protein tyrosine kinase activity, and biological process protein phosphorylation (Figure 3a). The most upregulated genes were associated with cellular component, membrane and membrane part. In addition, KOG (EuKaryotic Ortholog Groups) analysis indicated that 120 upregulated genes were related to signal transduction mechanisms (Figure 3b). KEGG (Kyoto encyclopedia of genes and genomes) analysis was also performed, and revealed that IRP-regulated genes are involved in plant−pathogen interaction, sphingolipid signaling, sphingolipid metabolism and phenylalanine metabolism (Figure 3c). These results suggested that IRP functions in regulating signal transduction at the cell membrane.

**Figure 3.**
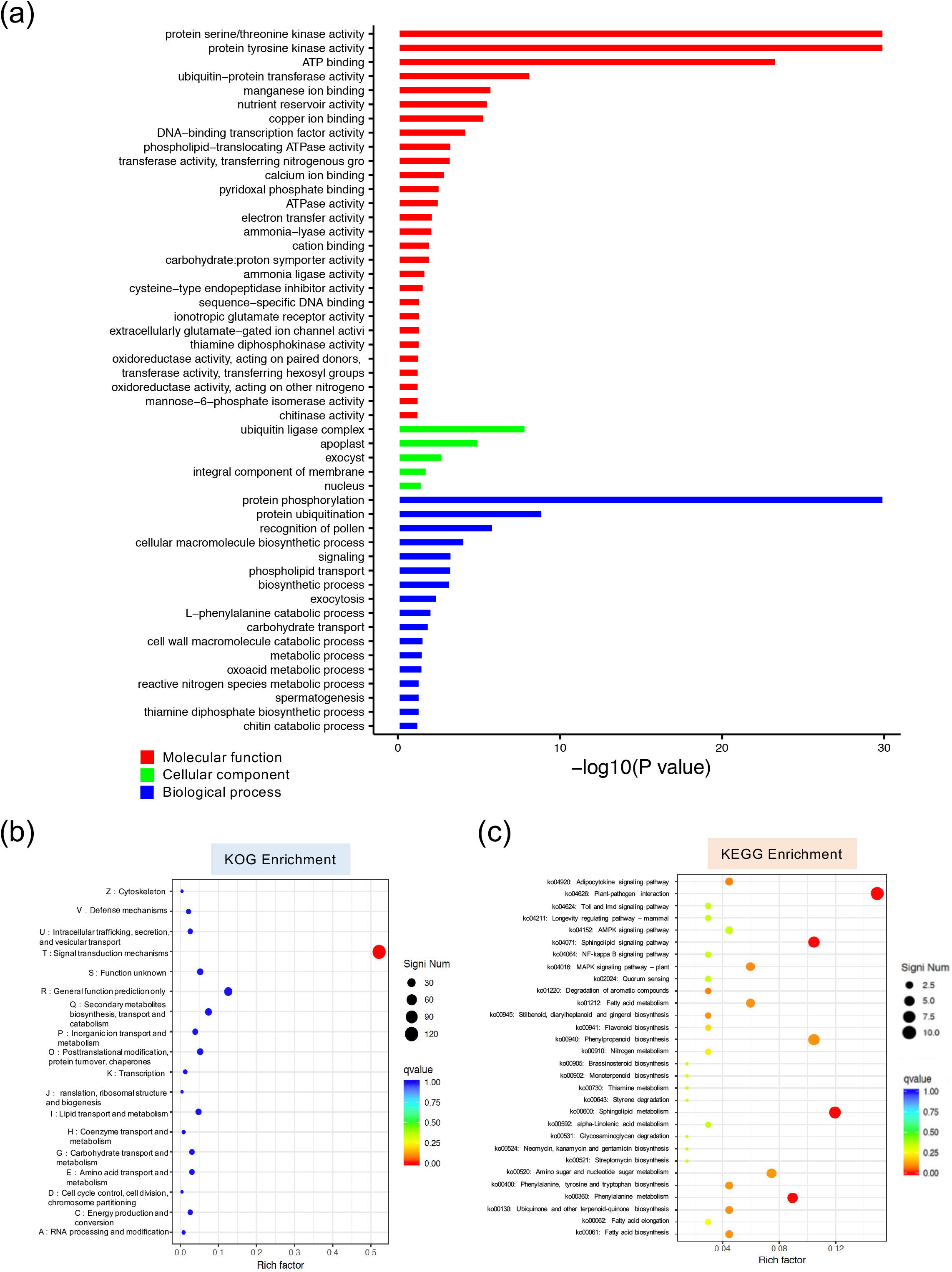
Functional analysis of genes upregulated by IRP. (a) Bar plots summarizing the gene ontology terms over-represented in upregulated genes by IRP. Bar color indicates the three categories of GO term: molecular functions (red), cellular components (green), and biological processes (blue). (b) Scatter plot of KOG pathway enrichment statistics. Rich factor is the ratio of IRP upregulated gene number annotated in this pathway term to all gene number annotated in this pathway term. Greater rich factor means greater intensiveness. q-value is corrected P-value ranging from 0 ~ 1, and its less value means greater intensiveness (indicated by the red dot). The dot size indicated the number of upregulated genes. The top 30 pathway terms enriched by KEGG database were displayed. (c) Scatter plot of KEGG pathway enrichment statistics. Rich factor is the ratio of IRP upregulated gene number annotated in this pathway term to all gene number annotated in this pathway term. Greater rich factor means greater intensiveness. q-value is corrected P-value ranging from 0 ~ 1, and its less value means greater intensiveness (indicated by the red dot). The dot size indicated the number of upregulated genes.

### IRP and chitin elicit a shared immune response

At the same time, we performed chitin treatment for RNA-seq analysis in order to compare the response regulated by chitin and IRP. An average of 55 million reads was obtained per sample with an average mapping rate of 91.8% to the rice genome (Table S2). In chitin treatment, a total of 1,222 genes were differentially expressed, consisting of 1,097 upregulated and 125 downregulated (Table S3). Interestingly, 84% of the DEGs (585) enhanced by IRP were also induced by chitin treatment (Figure 4a). GO results indicated that 68 GO terms are enriched in these common genes (Figure S2). Moreover, several DEGs common to IRP and chitin treatment belonged to GO terms categorized as cellular process and metabolic process, catalytic activity and binding, and membrane and membrane part (Table S4). Next, all DEGs identified in chitin and IRP treatments were subjected to hierarchical clustering analysis and divided into four clusters (Figure 4b). In cluster 1, the expression of 183 genes decreased in both IRP and chitin treatments. A total of 207 genes were highly induced by chitin in cluster 2, while they were not induced by IRP treatment. On the contrary, the expression level of 41 genes in cluster 3 was only enhanced in IRP treatment. In cluster 4, the expression level of 959 genes increased in both IRP and chitin treatments.

**Figure 4.**
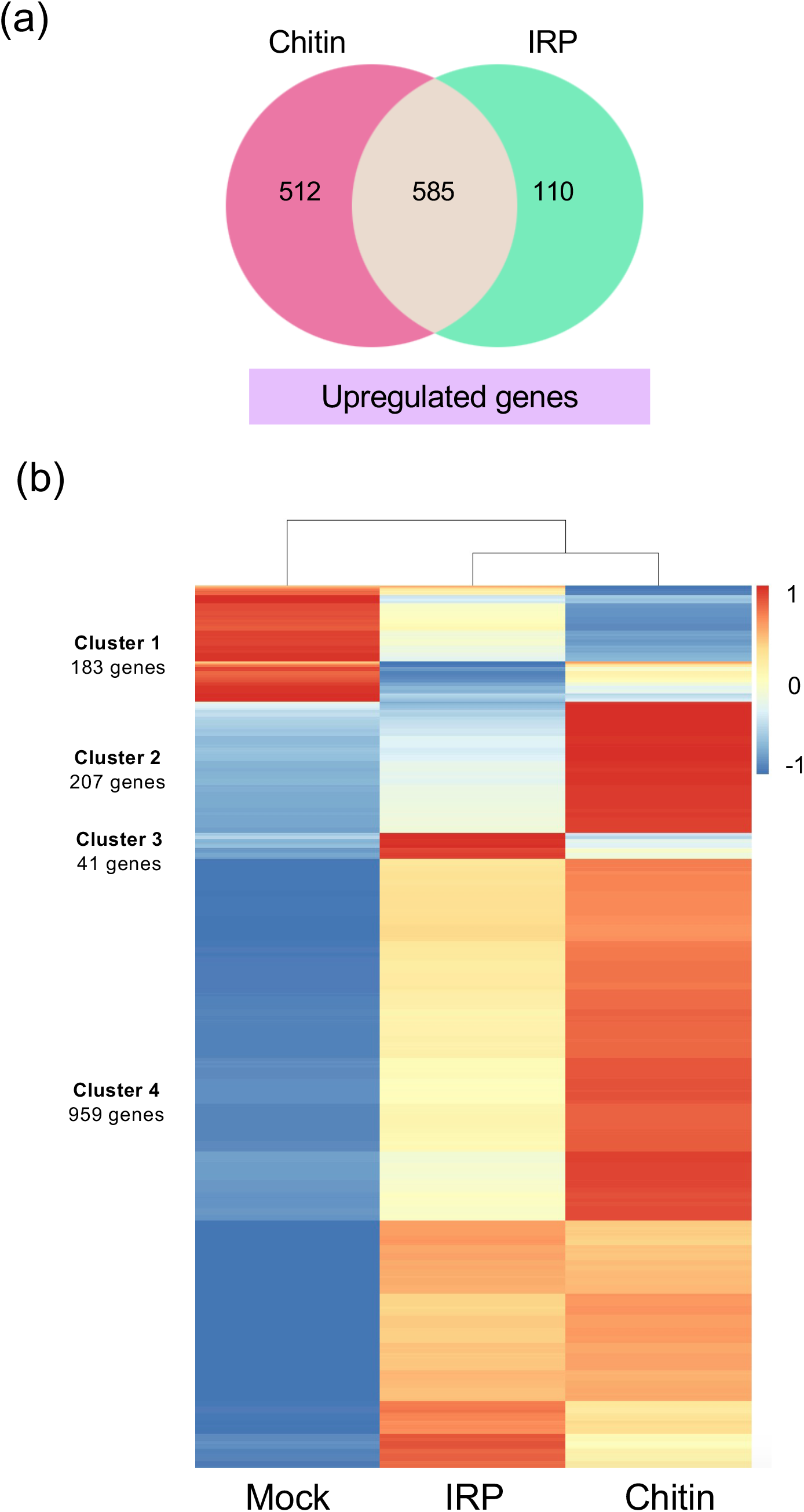
IRP and chitin trigger similar sets of genes. (a) Venn diagram showing the overlap in upregulated genes between chitin treatment and IRP treatment. (b) Clustering analysis on genes differentially expressed in response to chitin and IRP treatment. A total of 1,390 DEGs (Log_2_FC > 1, *p* < 0.05) from the RNA-seq were divided into four clusters using the default parameters of the clustering method in the R package pheatmap.

To further explore the relationship between IRP and chitin in rice immunity, we treated rice suspension cells with chitin and IRP together. The expression levels of both *IRP* and *PAL1* in cells treated with 500 nM IRP but without chitin were much lower than those in cells treated with 1 μg/mL chitin without IRP (Figure 5a, b). In the 1 μg/mL chitin condition, *IRP* and *PAL1* expression were further increased in an IRP concentration-dependent manner. In the presence of 2 μg/mL chitin, IRP treatment did not trigger statistically significant induction of *IRP* and *PAL1* expression. Next, we monitored the activation of two MAPKs, MPK3 and MPK6 (MPK3/6). Interestingly, the MPK3/6 activation level in the presence of 500 nM IRP was higher than that in the presence of 1 μg/mL chitin. IRP-dependent MPK3/6 activation was also observed in the presence of 2 μg/mL chitin (Figure 5c). It appears that IRP plays active roles in both gene expression and MAPK activation, but has a stronger effect on MAPK activation than on *IRP* and *PAL1* expression when compared with chitin.

**Figure 5.**
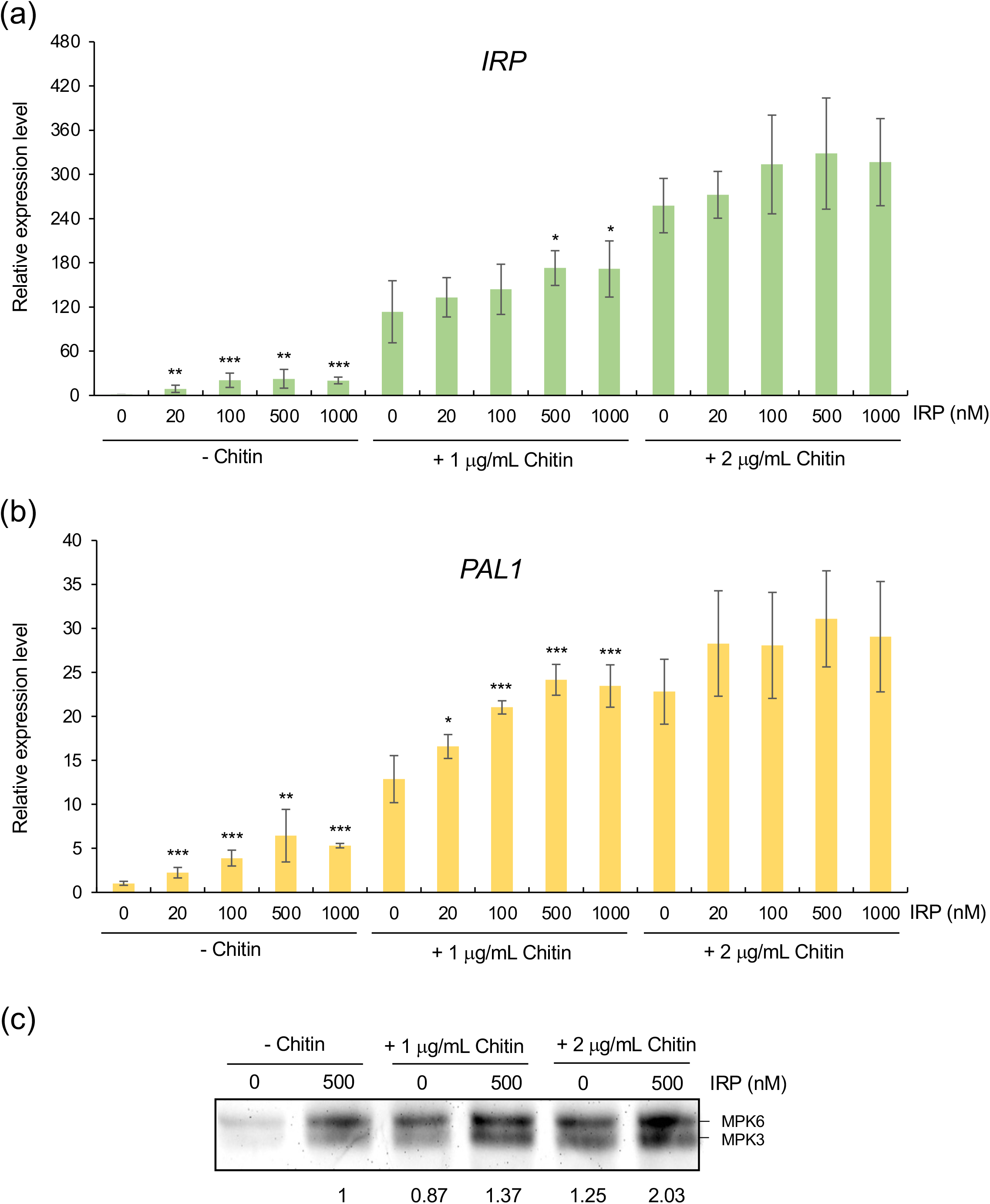
Chitin regulates IRP in rice immunity. (a, b) Relative expression of *IRP* (a) and *PAL1* (b) in rice suspension cells treated with different concentrations of IRP peptide and chitin for 1 h. Error bars indicate SD. Statistically significant differences between IRP treatment and mock IRP in the corresponding chitin condition are depicted with asterisks (*p < 0.05; **p < 0.01; ***p < 0.001; n=4, two-tailed *t*-test). (c) MAPK activation was detected after treatment for 15 min with different concentrations of IRP and chitin. The relative levels of MPK3/6 phosphorylation shown below the gel were determined with ImageJ software. Three independent experiments were performed and similar results were obtained.

### IRP and chitin regulate the same transcription factors

After sensing pathogen attacks, transcriptional reprogramming of thousands of genes is triggered by transcription (co)factors (Moore et al., 2011). In our RNA-seq results, several families of transcription factor such as WRKYs and APETELA2/ethylene response factor (AP2/ERF) were identified after IRP treatment (Table S1 and S3). We found that IRP and chitin promoted the expression of 14 and 18 *WRKY*s, respectively, and 13 of them overlapped in both treatments (Figure 6a, Table 1). Four *WRKY*s, *OsWRKY24*, *OsWRKY42*, *OsWRKY69*, and *OsWRKY70*, were selected for further analysis. Both IRP and chitin triggered the expression of these four *WRKY*s according to qPCR results (Figure 6b–e). In addition, IRP coupled with chitin treatment increased the expression level of the four *WRKY* genes compared with chitin treatment alone (Figure 6b–e). However, significant induction of the four selected *WRKY*s by IRP was no longer observed when chitin concentration reached 2 μg/mL. These results implied that chitin and IRP share the same pathway to control the expression of these *WRKY* genes.

**Figure 6.**
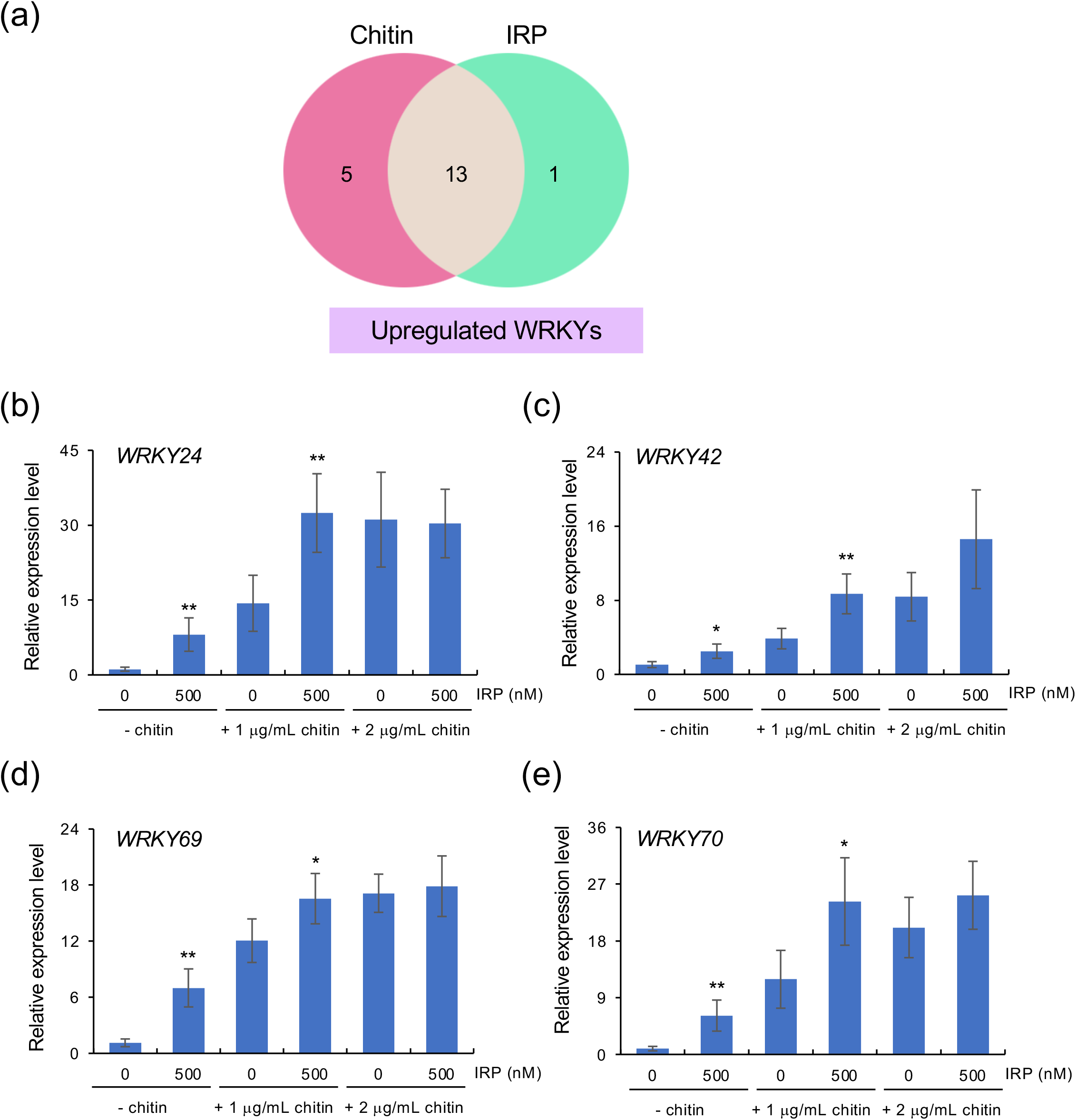
Chitin and IRP induce the expression of *OsWRKY*s. (a) Venn diagram showing the overlapping upregulation of *OsWRKY* transcription factors between chitin treatment and IRP treatment. (b–e) The expression levels of *OsWRKY24* (b), *OsWRKY42* (c), *OsWRKY69* (d), and *OsWRKY70* (e) were analyzed by qPCR in rice suspension cells treated with the indicated concentration of chitin and IRP for 1 h. Error bars indicate SD. Statistically significant differences between IRP treatment and mock IRP in the corresponding chitin condition are depicted with asterisks (*p < 0.05; **p < 0.01, n=4, two-tailed *t*-test).

**Table 1.** WRKY transcription factors identified in chitin and IRP treatments. Y, identified; N, unidentified.

| Gene ID | Description | Chitin treatment | IRP treatment |
| --- | --- | --- | --- |
| LOC_Os01g09080 | WKRY107 | Y | N |
| LOC_Os01g09100 | WKRY10 | Y | N |
| LOC_Os01g46800 | WKRY15 | Y | Y |
| LOC_Os01g60540 | WKRY20 | Y | Y |
| LOC_Os01g60600 | WKRY108 | Y | N |
| LOC_Os01g60640 | WKRY21 | Y | Y |
| LOC_Os01g61080 | WKRY24 | Y | Y |
| LOC_Os02g08440 | WKRY71 | Y | Y |
| LOC_Os02g26430 | WKRY42 | Y | Y |
| LOC_Os02g53100 | WKRY32 | Y | Y |
| LOC_Os05g27730 | WKRY53 | Y | Y |
| LOC_Os05g39720 | WKRY70 | Y | Y |
| LOC_Os05g49100 | WKRY49 | Y | N |
| LOC_Os05g49210 | WKRY43 | Y | Y |
| LOC_Os05g50700 | WKRY111 | Y | N |
| LOC_Os06g05380 | WKRY73 | N | Y |
| LOC_Os06g06360 | WKRY113 | Y | Y |
| LOC_Os06g44010 | WKRY28 | Y | Y |
| LOC_Os08g29660 | WKRY69 | Y | Y |

Y, identified; N, unidentified

### ABA enhances*IRP* expression

We found that IRP elevated *OsWRKY24* expression, which is also known to be upregulated by abscisic acid (ABA) treatment (Zhang et al., 2009) (Figure 6b). Moreover, RNA-seq results revealed that an ABA-inducible BHLH-type transcription factor was also induced by IRP (Table S1). These results raise the possibility that IRP is involved in ABA-regulated signaling pathways. Therefore, the expression level of *IRP* after ABA treatment was tested. The *OsHOX22* gene, which functions in ABA biosynthesis, was used as a marker of activation of ABA signaling (Liu et al., 2012). We treated suspension cells with ABA for 1.5 h and found that the expression of *IRP* was indeed upregulated by ABA (Figure 7).

**Figure 7.**
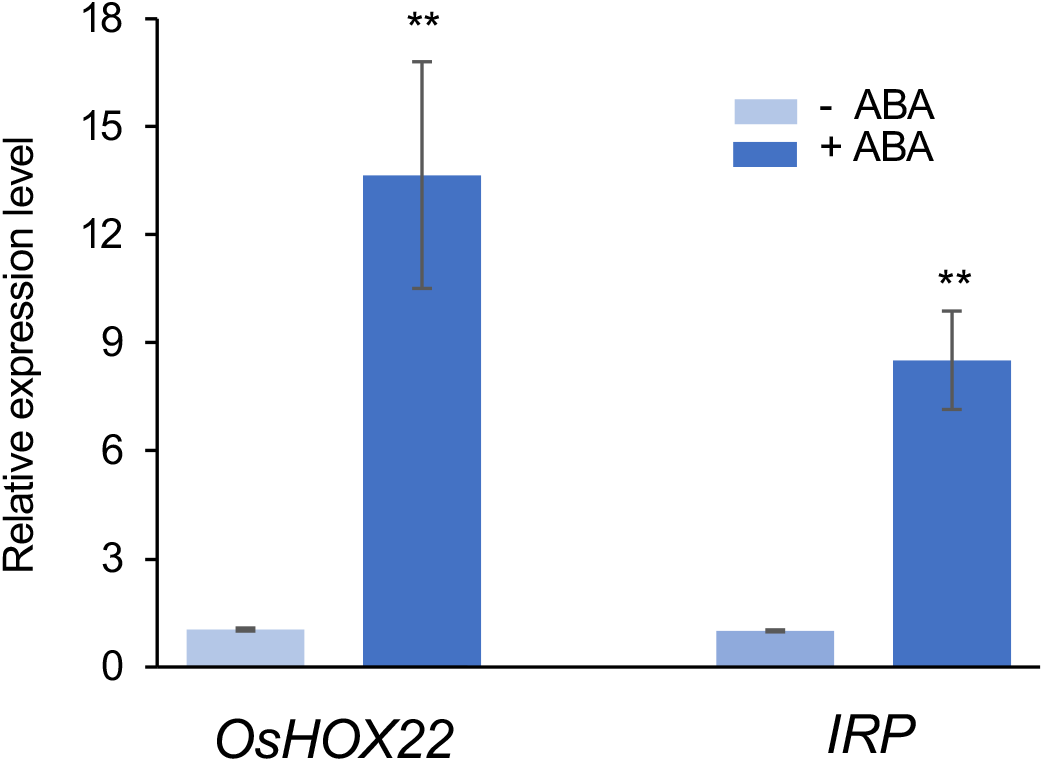
ABA induces *IRP* expression. Rice suspension cells were treated with 100 nM ABA for 1.5 h and the relative expression levels of *OsHox22* and *IRP* were detected by qPCR. Error bars indicate SD. Statistically significant differences between treatment and mock are depicted with asterisks (**p < 0.01, n=3, two-tailed *t*-test).

## DISCUSSION

Since the first plant peptide, systemin, was identified in tomato, many peptides have been isolated from plants (Pearce et al., 1991). Recently, the roles of peptides as signaling molecules have been recognized not only in growth and development but also in biotic and abiotic stress responses (Olsson et al., 2019). However, despite its importance as a crop, our knowledge of peptides in rice is limited. The first studied peptide in rice, *Os*PSK, was found to play a critical role in cell proliferation (Yang et al., 1999). Later, several database searches for *Arabidopsis* homologs have resulted in the identification of CLAVATA3 (CLV3)/EMBRYO SURROUNDING REGION (ESR) (CLE), RALF, and PSK families in rice (Sawa et al., 2008; Sauter, 2015; Sharma et al., 2016). Recently, we identified five *Os*RALFs and *Os*PSK4 as potential DAMPs in rice immunity by combining transcriptomics with proteomics strategies (Wang et al., 2020). Moreover, we also isolated the novel peptide family IRP, whose homologs exist only in Poaceae, and detected a 20-aa C-terminal portion of an IRP peptide in rice suspension cell medium whose concentration was increased by chitin. In this study, we synthesized the C-terminus of this IRP peptide without the N-terminal signal peptide sequence and tested its function. The expression of *IRP* itself was induced in suspension cells treated with IRP peptide, which is a typical property of hormone-like peptides (Huffaker et al., 2006; Chen et al., 2014; Hou et al., 2014).

After perceiving PAMPs, plants are able to produce DAMPs to amplify immune responses (Yamaguchi and Huffaker, 2011). DAMPs include oligomeric fragments of cuticles and plant cell walls, proteins, peptides, nucleotides, and amino acids (Choi and Klessig, 2016). In the past two decades, some peptide families have been found to function as DAMPs, such as systemin, Pep, RALF, and PIP. However, the number of peptide DAMP families is expected to be much higher as thousands of small protein-encoding genes exist in the *Arabidopsis* as well as the rice genome (Pan et al., 2013; Takahashi et al., 2019). In general, peptide DAMPs have several common features: (1) PAMP treatment increases transcription of DAMP precursors; (2) DAMPs and PAMPs activate similar immune responses; and (3) DAMP receptor(s) are required for full activation of PTI signaling and resistance against pathogen infection (Yamaguchi et al., 2010; Tintor et al., 2013; Hou et al., 2014). In our previous work, the transcription of *IRP* was induced by the PAMP chitin as well as by PGN (Wang et al., 2020). In this study, overexpression of *IRP* enhanced the resistance of rice plants to a fungal pathogen. IRP peptide not only induced the expression of the defense gene *PAL1*, but also enhanced MAPK activation. Intriguingly, 84% of IRP-upregulated genes overlapped with genes induced by chitin, implying that IRP works as a DAMP to amplify the chitin signaling pathway and IRP is the first studied peptide DAMP in rice immunity (Figure 4).

Besides, a total of 14 and 18 *OsWRKY* transcription factors were induced by IRP and chitin, respectively, 13 of which overlapped in both treatments, strongly suggesting that IRP shares many of the same WRKY signaling components with chitin to regulate the immune response. Among the common 13 WRKYs, we focused on four WRKYs, *Os*WRKY24, *Os*WRKY42, *Os*WRKY69, and *Os*WRKY70*. Os*WRKY24 is a homolog of *Os*WRKY70, which belongs to group I-type WRKYs. It has been reported that an *MPK6* RNAi mutant suppresses the expression level of *OsWRKY24* and *OsWRKY70.* OsWRKY70 is a substrate of MPK3/6 (Li et al., 2015). Overexpression of *OsWRKY70* prioritizes defense over growth by positively regulating JA and negatively regulating gibberellin (GA), resulting in the enhancement of resistance against the chewing herbivore *Chilo suppressalis*. Together, these results raise the possibility that IRP regulates *Os*WRKY24 and *Os*WRKY70 through MPK3/6. In addition, both *IRP* and *OsWRKY24* were induced by ABA, suggesting that IRP also contributes to the ABA signaling pathway (Figure 7). A previous study indicated that *OsWRKY69* is induced by blast fungus (Bagnaresi et al., 2012), while *Os*WRKY42 plays a negative role in rice resistance to blast fungus (Cheng et al., 2015). Further analyses of how IRP controls these negative WRKYs will be an interesting task in rice immunity.

In this study, we found that most of the DEGs regulated by IRP were also observed in chitin treatment, implying that IRP is a critical DAMP, working downstream of chitin. On the other hand, no enhancement of ROS production was detected in suspension cells treated with IRP, indicating that IRP cannot replicate all of the chitin-induced immune responses (Figure 2h, i). In addition to IRP, we previously isolated another 235 small secreted proteins induced by the rice blast fungus *M. oryza*e and its elicitor chitin, including six known DAMP family proteins: OsRALF6, OsRALF7, OsRALF8, OsRALF26, OsRALF31, and OsPSK4 (Wang et al., 2020). Similarly, previous analysis of flg22- and elf18-induced transcription led to the identification of many DAMPs, *PSK4*, *PSY1*, *IDA*, and *PIP*s (Hou et al., 2014). This implies that plants secrete several DAMPs, simultaneously to orchestrate immune responses. Interestingly, there are no visible effects on resistance to rice blast fungus in *IRP* knockout plants (Figure S1), perhaps due to compensation by other DAMPs. In addition to fungal chitin, the expression of *IRP* is also enhanced by a bacterial elicitor, PGN (Wang et al., 2020). It is therefore possible that IRP is involved in immune responses regulated by other PAMPs. In summary, we propose a working model for IRP (Figure 8). Plants recognize pathogens and then induce several DAMPs, including IRP and others, which are secreted into the apoplast and sensed by their corresponding receptors. IRP activates the MAPK cascade and then regulates WRKY transcription factors; these in turn induce defense genes, leading to amplification of the rice immune response. Therefore, to uncover IRP function, one of the most crucial goals is to identify IRP receptor(s) and we will address this important question in the future.

**Figure 8.**
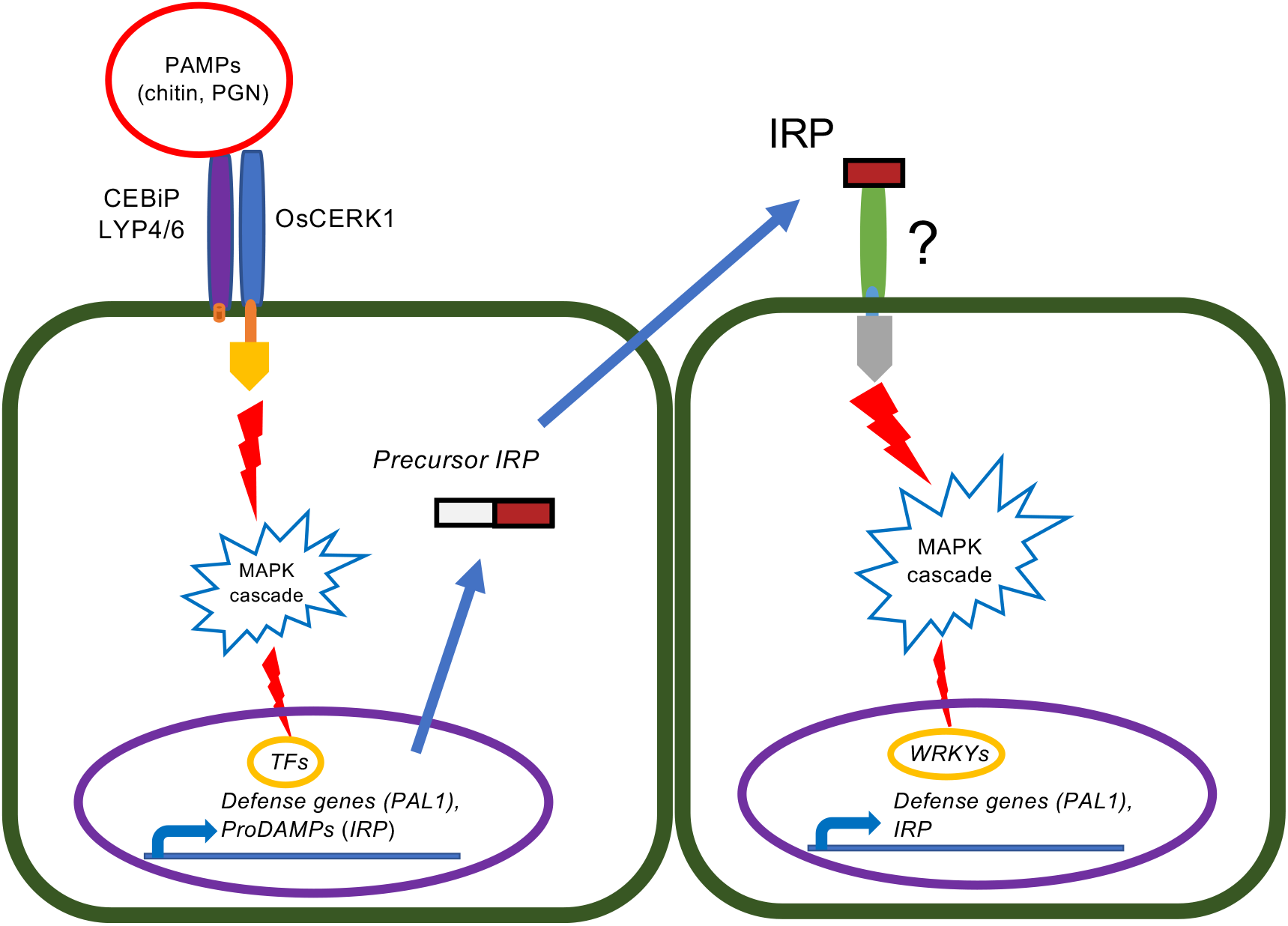
Model of IRP as a DAMP in rice PTI. (a) Different PAMPs, chitin and PGN, are perceived by PRRs and activate MAPKs and transcription factors (TFs). TFs induce the expression of defense genes, such as *PAL1*, and genes encoding precursors of peptide DAMPs, including *IRP*. The IRP precursor is cleaved during secretion into the apoplast of the mature form of IPR peptide; It is recognized by its receptor, triggering further PTI responses including MAPK activation and TF expression.

## ACKNOWLEDGEMENTS

We thank the members of the Laboratory of Signal Transduction and Immunity at PSC, and the Plant Immune Design Group at Okayama University, for invaluable support and discussions. We thank Dr. Jian-Kang Zhu (Shanghai Center for Plant Stress Biology) for providing us the CRISPR-Cas9 vector. This work was supported by the Chinese Academy of Sciences, Shanghai Institutes for Biological Sciences, Shanghai Center for Plant Stress Biology, CAS Center of Excellence for Molecular Plant Sciences, Strategic Priority Research Program of the Chinese Academy of Sciences (XDB27040202), the Chinese Academy of Sciences Hundred Talents Program (173176001000162114), the National Natural Science Foundation of China (31572073 and 31772246), JSPS KAKENHI (26450055, 17K07668, and 20H02988), and the Ohara Foundation.

## MATERIALS AND METHODS

### Plant growth and suspension cells

Rice plants (*Oryza sativa* L. ssp. *japonica* cv. Nipponbare) were grown in a greenhouse under a 16 h light/8 h dark cycle at 28°C/22°C. *IRP* was cloned to into the p2K1 vector for overexpression experiments (Wang et al., 2020). Two specific single guide RNAs (sgRNAs) were designed for CRISPR/Cas9 mutants (Table S5) and were introduced individually into the Cas9 vector pCBSG03 (Lu et al., 2017). The final constructs were transformed into Nipponbare calli via *Agrobacterium* strain EHA105. Rice suspension cells derived from Nipponbare calli were grown in R2 liquid medium at 30°C and the medium was changed once a week (Hayakawa et al., 1992, Wong et al., 2018).

### Rice plant infection and suspension cell treatment

For the infection assay, four-week-old rice plants were punched with *M. oryzae* (race 007.0) and covered by glass boxes to maintain high humidity. After one week, leaves were collected and lesion length was measured (Kawano et al., 2010). Chitin (hexa-o-acetylchitoheptaose, Carbosynth) was dissolved in H_2_O at a concentration of 10 mg/ml as a stock. IRP peptide was synthesized by Shanghai Top-peptide Biochemical Company and dissolved in H_2_O at a concentration of 2 mM as a stock. Rice suspension cells were subcultured for 12 h in fresh R2 liquid medium before treatments.

### Quantitative real-time PCR

Plant and suspension cell samples were harvested, frozen in liquid nitrogen and stored at −80°C. Total RNA was extracted by TRIzol reagent (Invitrogen), followed by reverse transcription using a cDNA synthesis kit (Vazyme) according to the manufacturer’s protocol. Quantitative real-time reactions were performed on a 6500 Real-Time PCR System (Applied Biosystems) using ChamQ SYBR qPCR Master mix (Vazyme). Relative expression level values were calculated using the 2^−ΔΔCT^ method (Livak and Schmittgen, 2001). To normalize gene expression across different samples, rice *Ubiquitin* (*LOC_Os03g13170.1*) was used as an internal control. Primers are listed in Table S5.

### RNA-seq assay

RNA quality was checked and quantity measured by Qubit2.0 RNA kit (Life Technologies) to equalize the quantity of each sample. The VAHTS mRNA-seq V2 Library Prep Kit for Illumina was used to construct transcriptome libraries, which were sequenced on an Illumina HiSeq XTen sequencing platform (Sangon Biotech). After filtering low-quality reads by Trimmomatic, clean reads were mapped to the rice genome (MSU7) using HISAT2. Gene expression was calculated as TPM (transcripts per million). FDR< 0.05 and at log_2_FC > 1 were used as the significance cut-off for genes differentially expressed in treated rice suspension cell samples compared to the corresponding mock samples.

The k-means clustering method was used to cluster genes up-regulated by IRP treatment or chitin in suspension cells (k = 6), using the Pearson correlation coefficient as the distance metric. A heat map was generated based on the TPM after log_2_ transformation, using the R package Pheatmap based on the following parameters: clustering method = “complete” and cluster_row = “TRUE”. Genes upregulated by IRP were subjected to GO (gene ontology), KGO (EuKaryotic Ortholog Groups) and KEGG (Kyoto Encyclopedia of Genes and Genomes) enrichment analysis.

### Detection of MAPK phosphorylation

Treated rice suspension cells were ground in liquid nitrogen and suspended in extraction buffer (50 mM HEPES (pH 7.4), 5 mM EDTA, 5 mM EGTA, 50 mM β-glycerophosphate, 10 mM Na_3_VO_4_, 10 mM NaF, 2 mM DTT, cOmplete EDTA-free protease inhibitor cocktail (Roche)) (Wang et al., 2020). Protein concentration was determined by Pierce 660 nm Protein Assay (Thermo Fisher Scientific). Immunoblot of phosphorylated MAPKs was performed using anti-phospho p44/42 MAPK antibody as the primary antibody and peroxidase-conjugated goat anti-rabbit IgG (Sigma) as the secondary antibody.

### ROS measurement

Rice suspension cells were collected in 96-well plates and incubated at 30°C for 16 h. For treatment, the medium was replaced by fresh liquid medium with 0.5 μM L-012 (Sigma-Aldrich), 10 μg/mL horseradish peroxidase (HRP, Sigma-Aldrich) and 2 μg/mL chitin or 1 μM IRP. Chemiluminescence was monitored using a luminescence microplate reader (Thermo Fisher Scientific).

## SUPPLEMENTAL FIGURES

**Figure 1.**
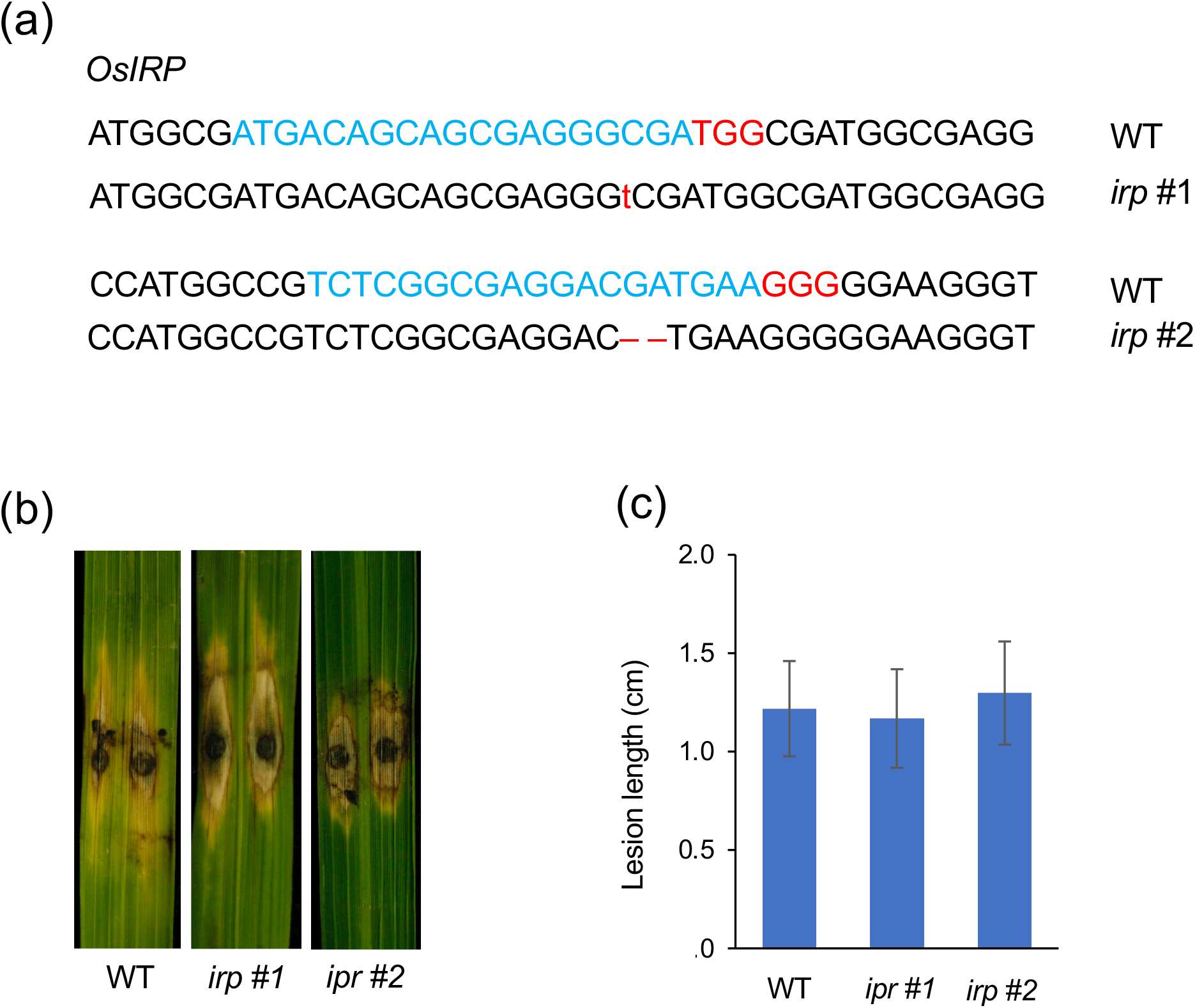
*IRP* mutants show no visible phenotype after blast fungus infection. (a) The two sgRNA:Cas9 targets and corresponding PAMs (protospacer-adjacent motifs) of *IRP* are shown in blue and red, respectively. The mutation of transgenic plants was confirmed by PCR. Deletions and an insertion are marked in red. (b) The phenotype of *IRP* mutants after blast fungus infection. (c) Quantitative analysis of lesion size in the rice plants one week after blast fungus infection.

**Figure 2.**
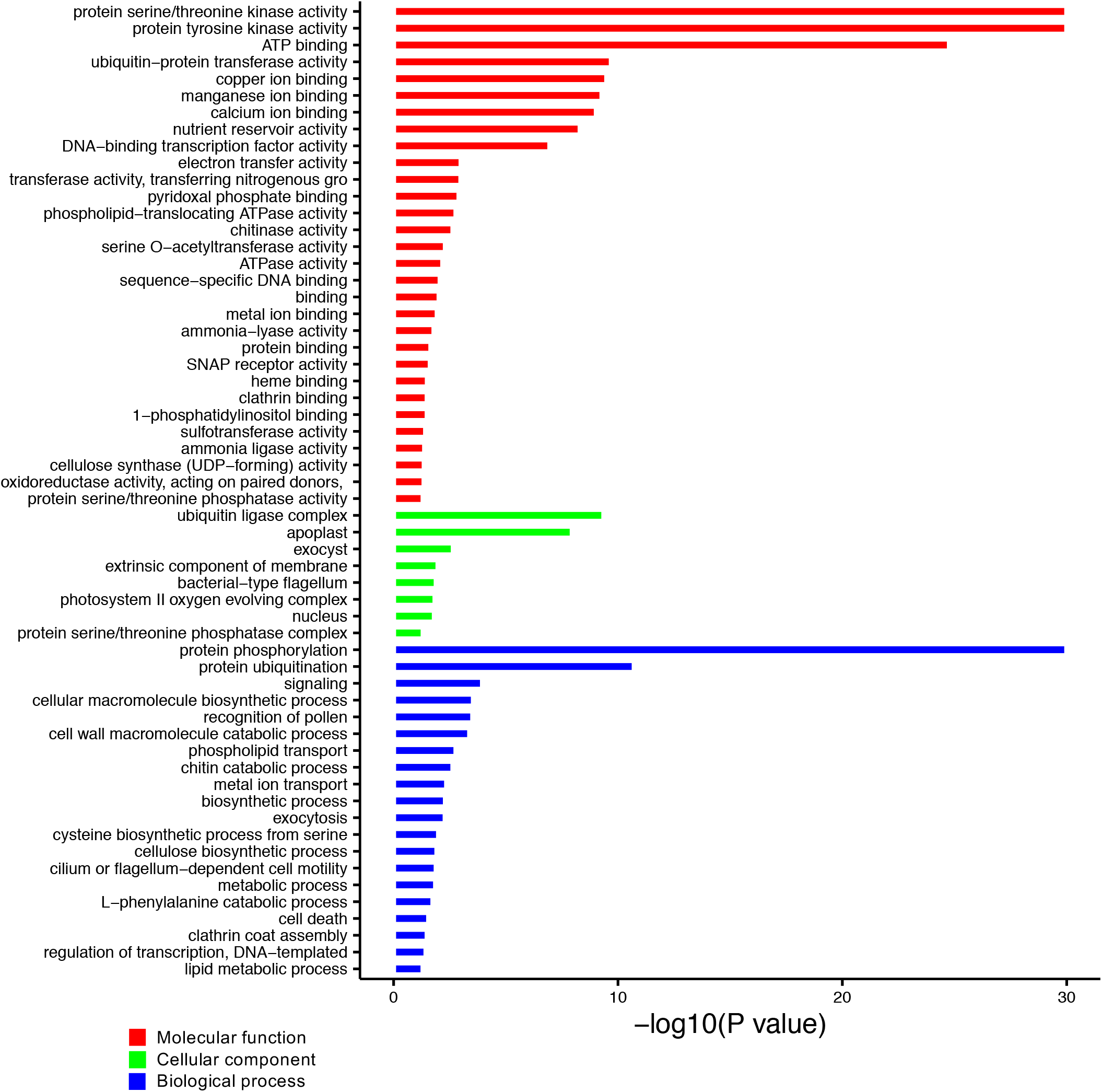
GO analysis of overlapping genes upregulated by IRP and chitin. Bar color indicates the three categories of GO term: molecular functions (red), cellular components (green), and biological processes (blue).

**Supplimental Table 1.**
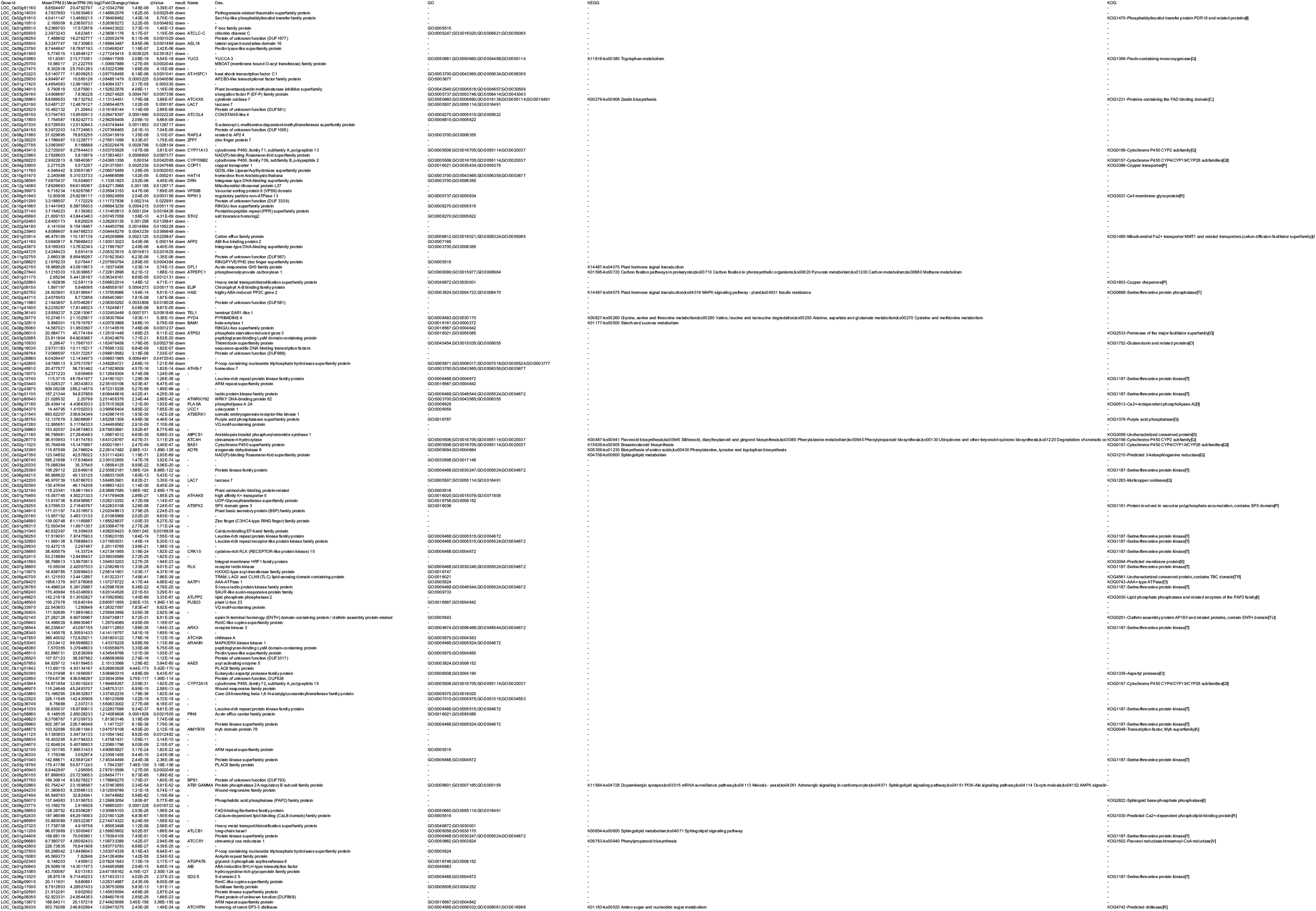
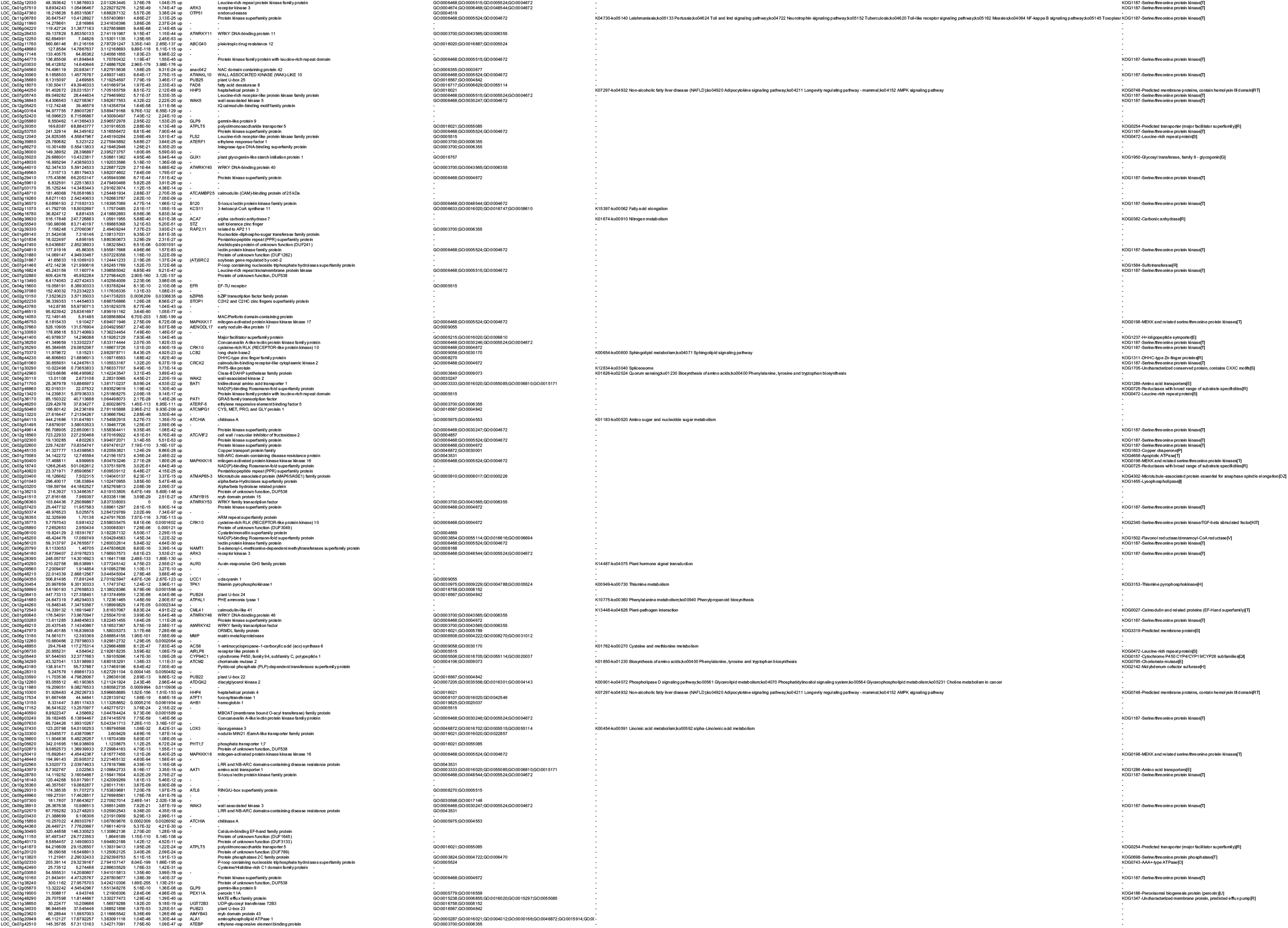
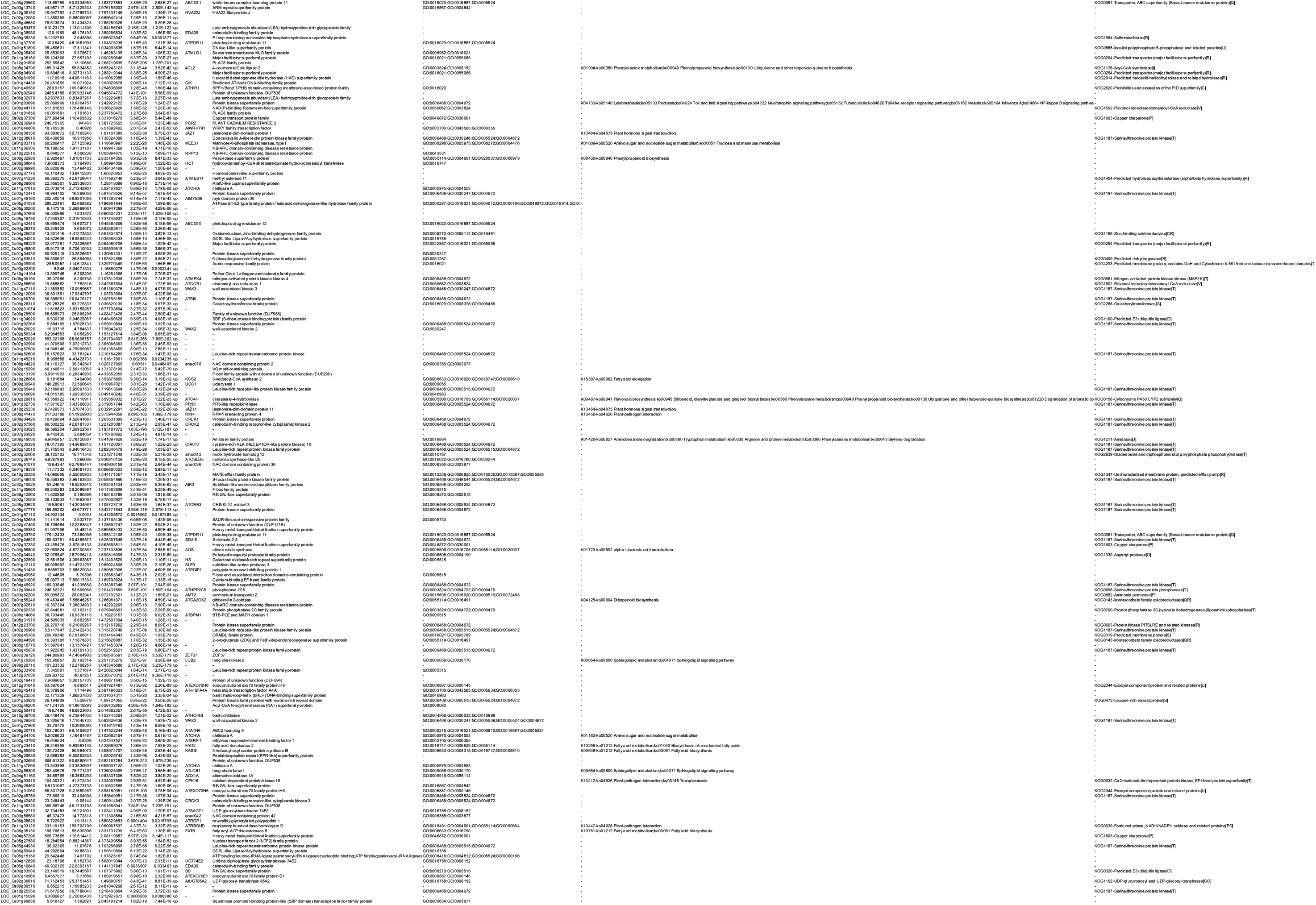
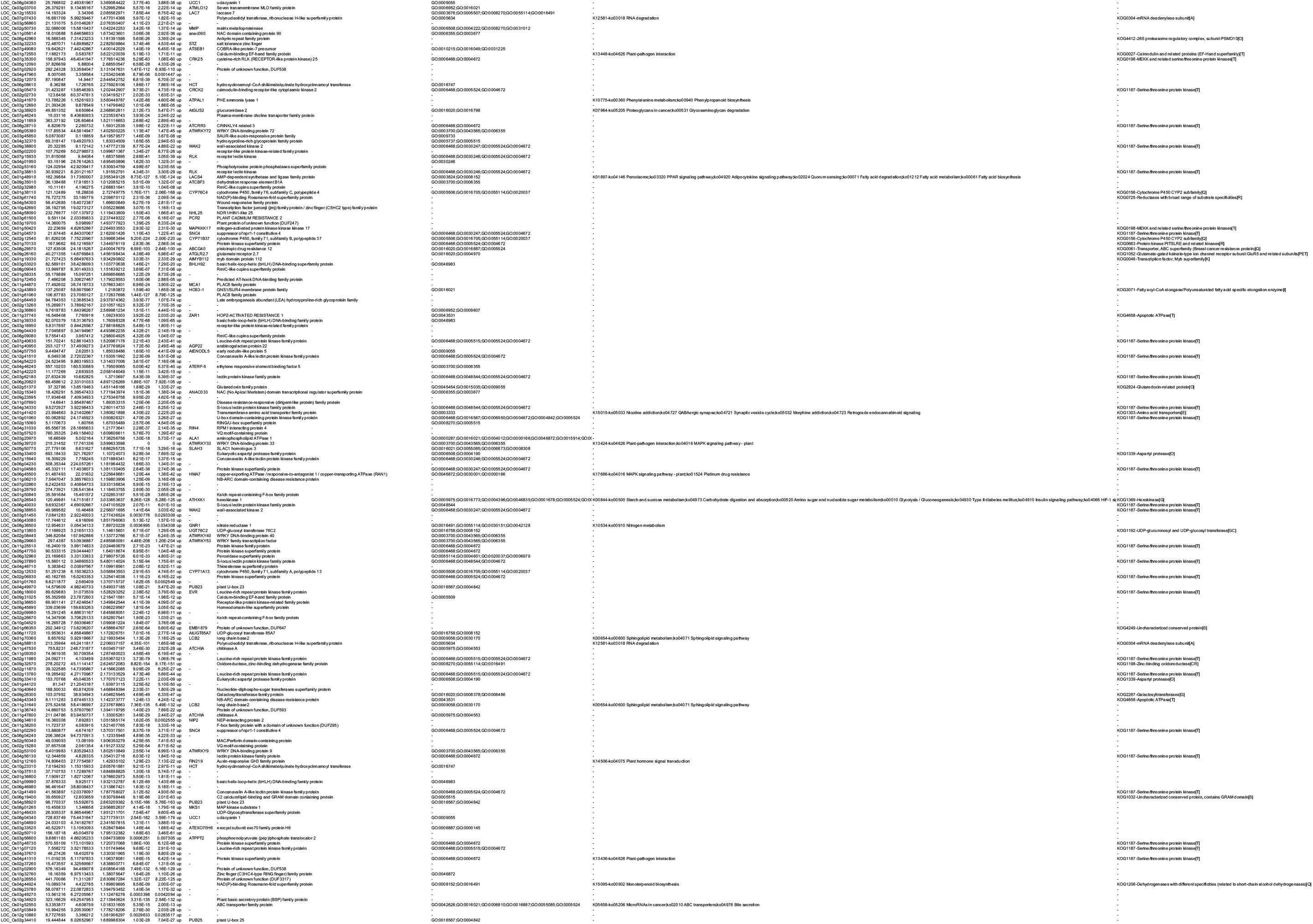
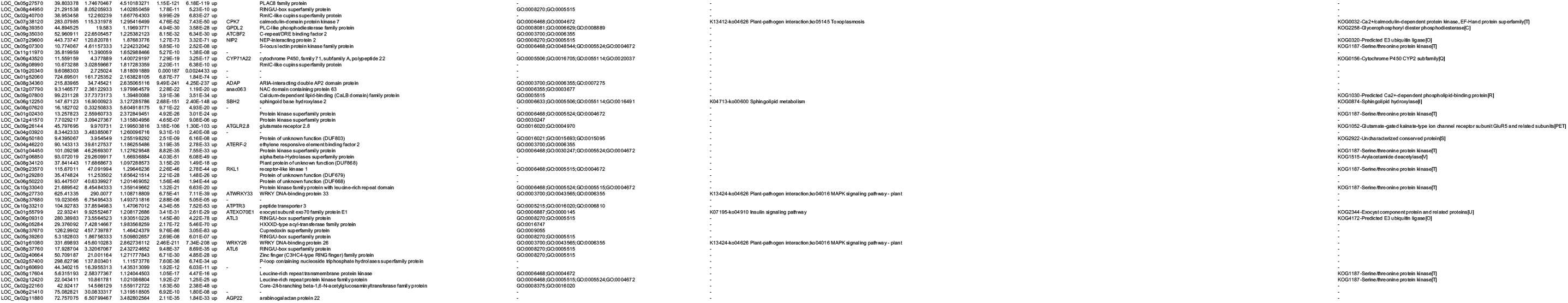
DEGs identified by IRP treatment.

**Supplimental Table 2.**
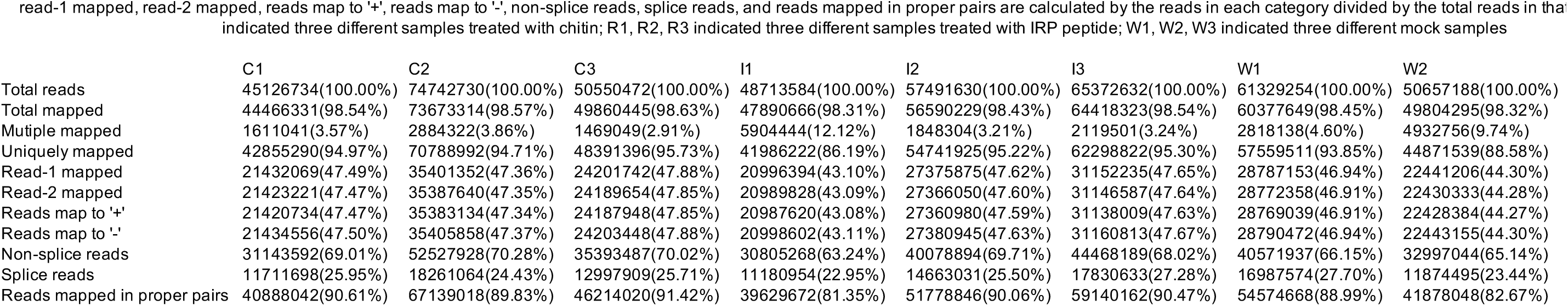
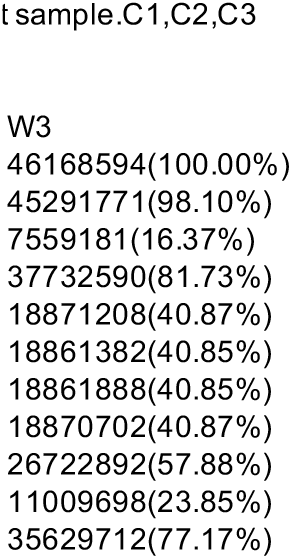
Mapping stastics.The percentage in total mapped,mutiple mapped, uniquely mapped,read-1 mapped, read-2 mapped, reads map to ‘+’, reads map to ‘−’, non-splice reads, splice reads, and reads mapped in proper pairs are calculated by the reads in each category divided by the total reads in tha indicated three different samples treated with chitin; R1, R2, R3 indicated three different samples treated with IRP peptide; W1, W2, W3 indicated three different mock samples

**Supplimental Table 3.**
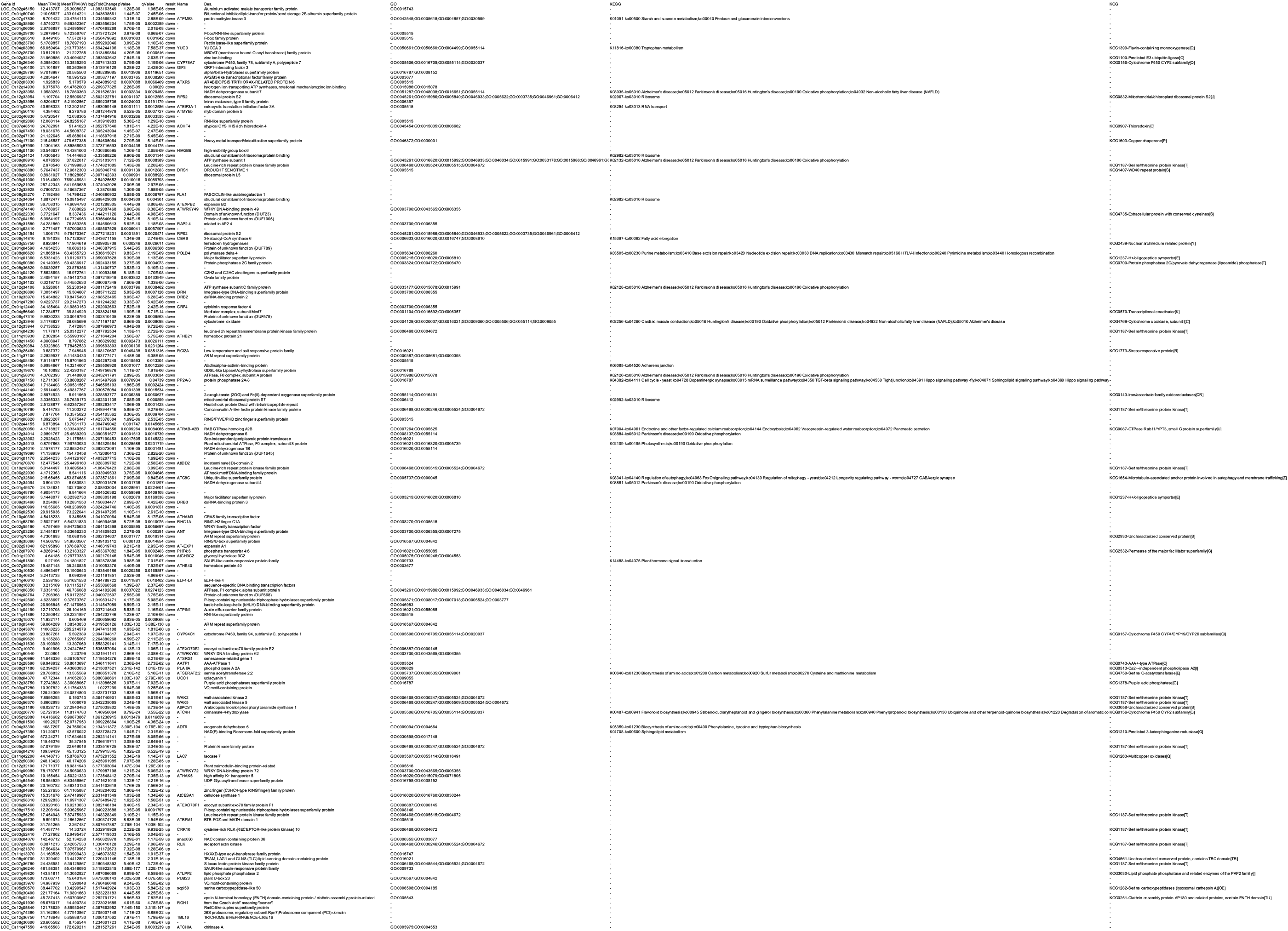
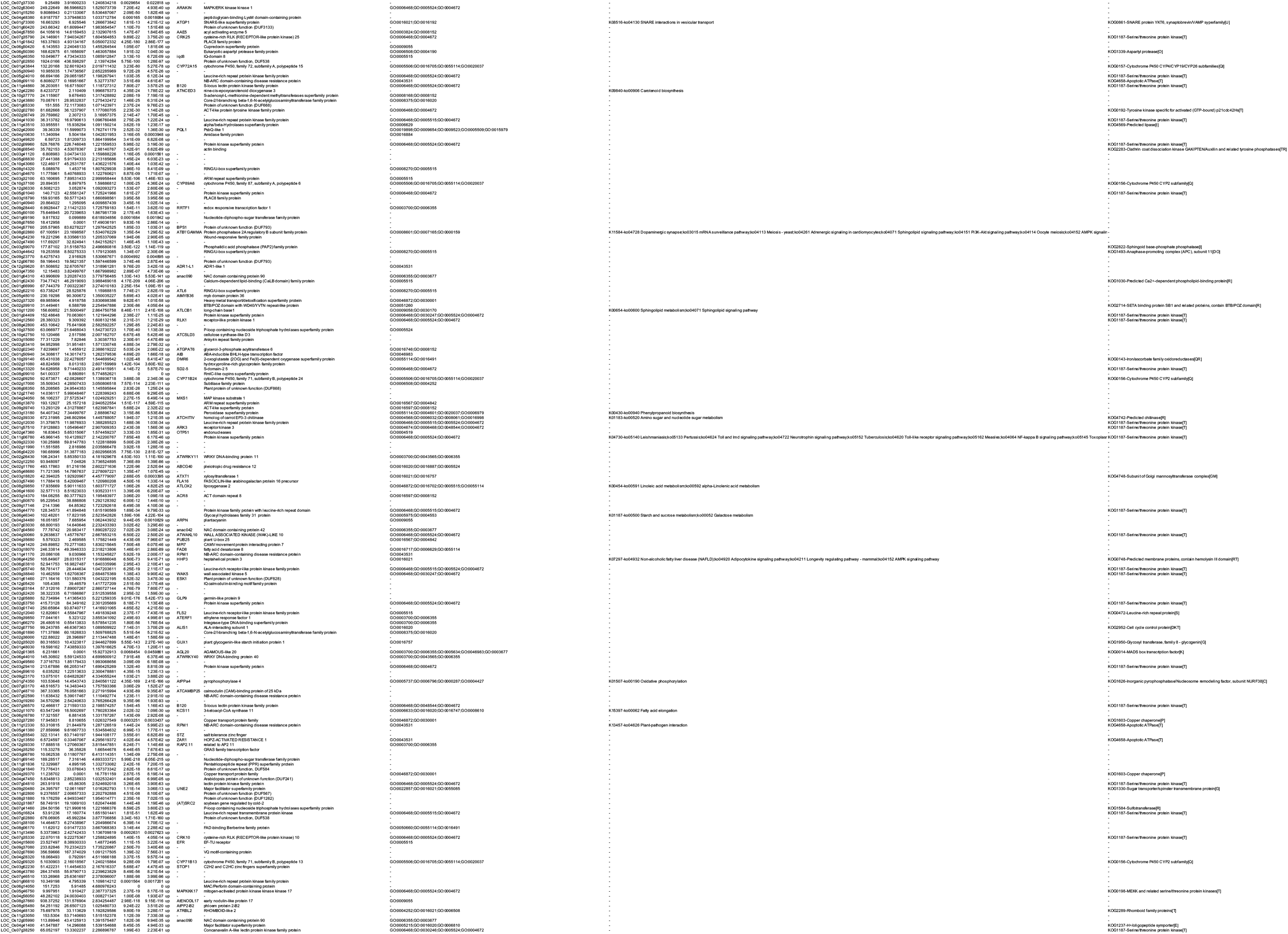
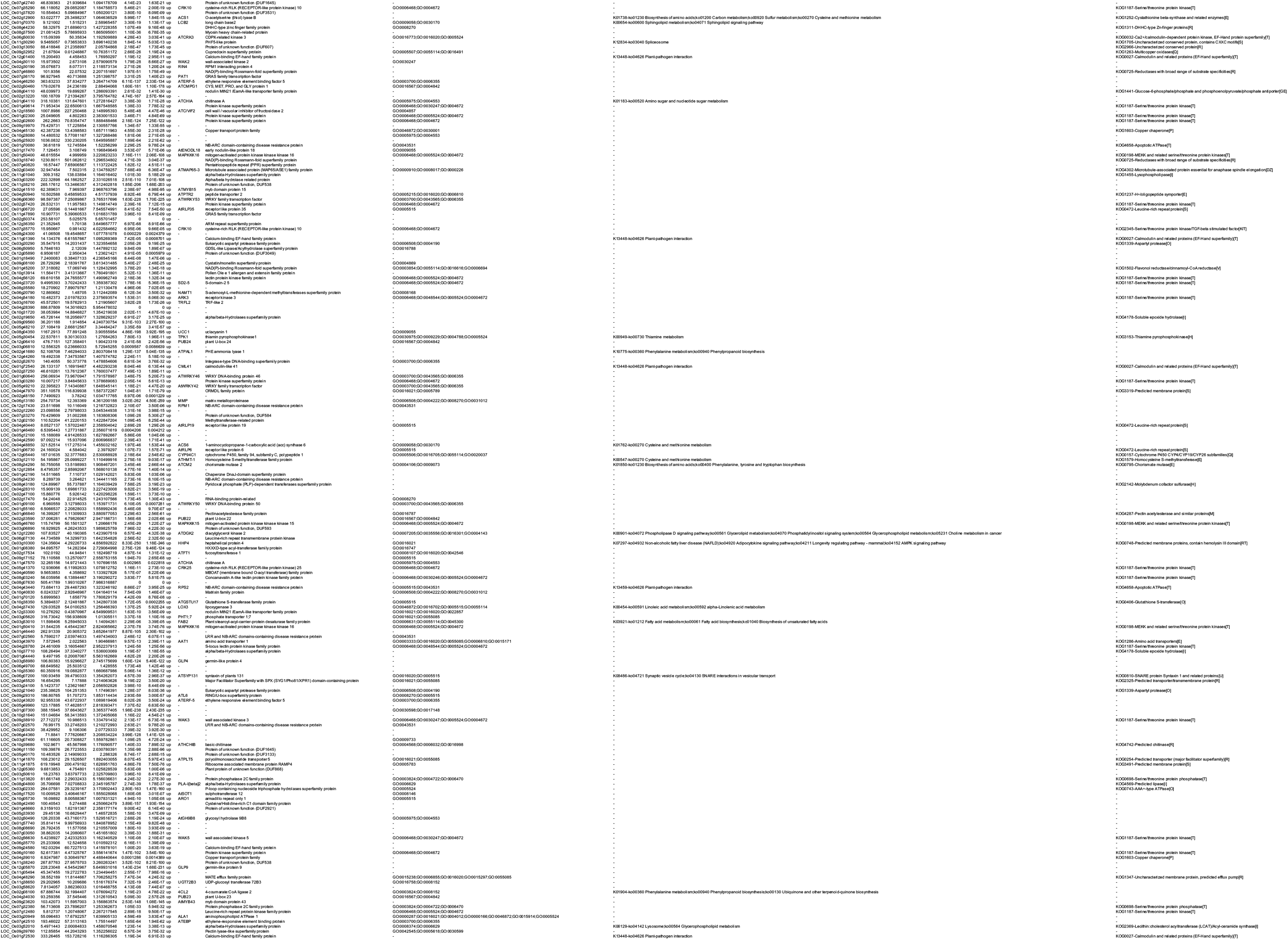
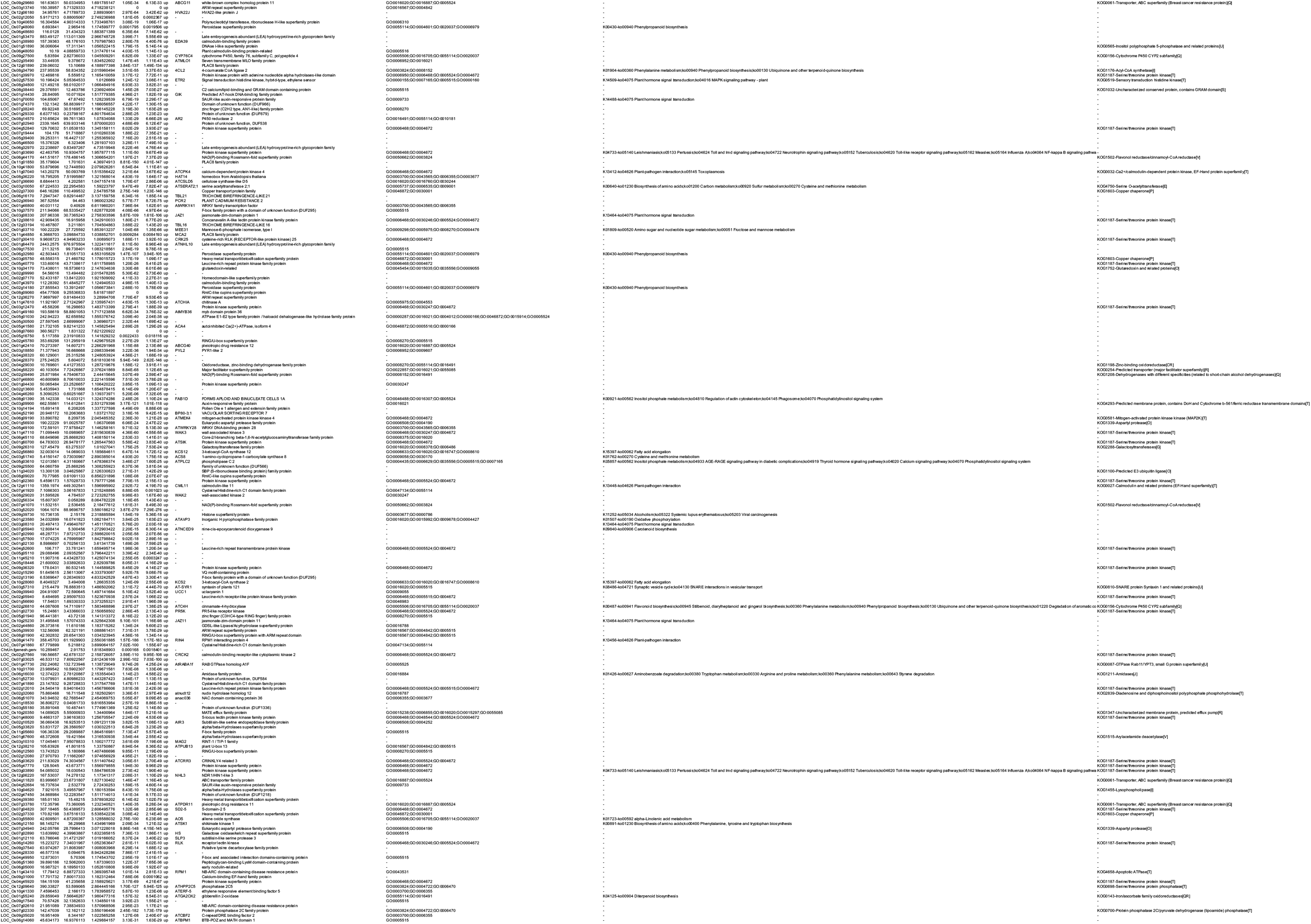
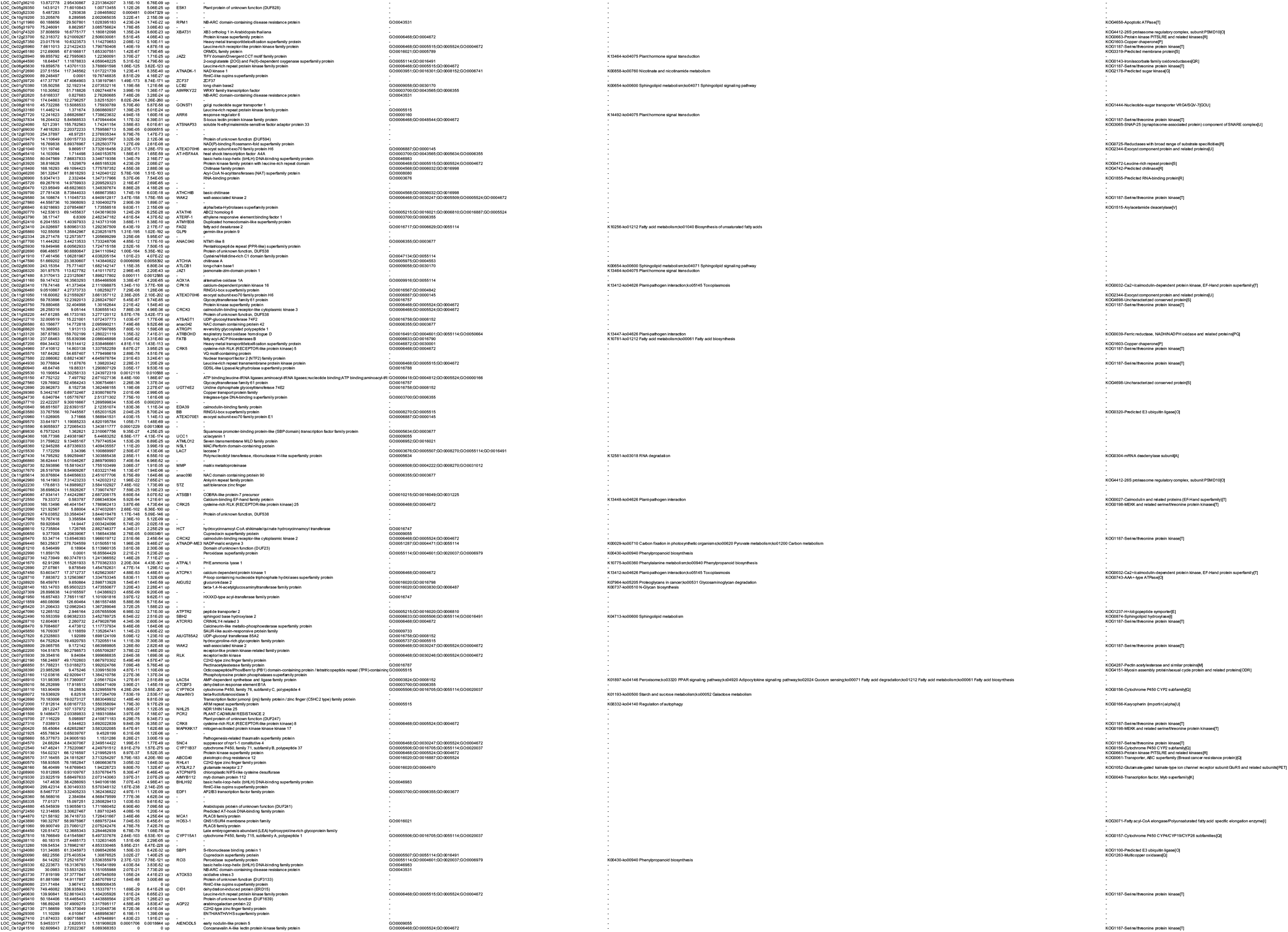
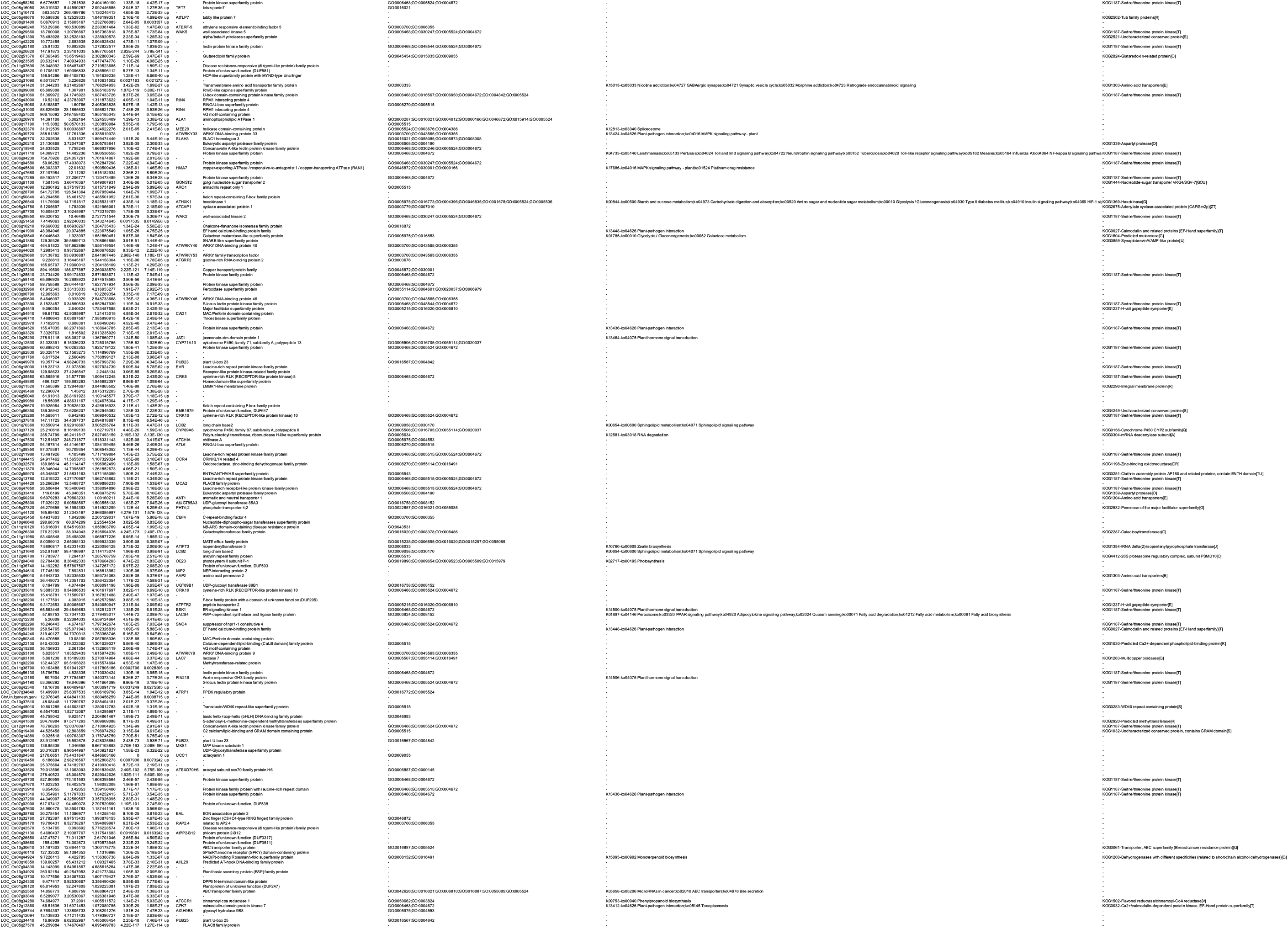
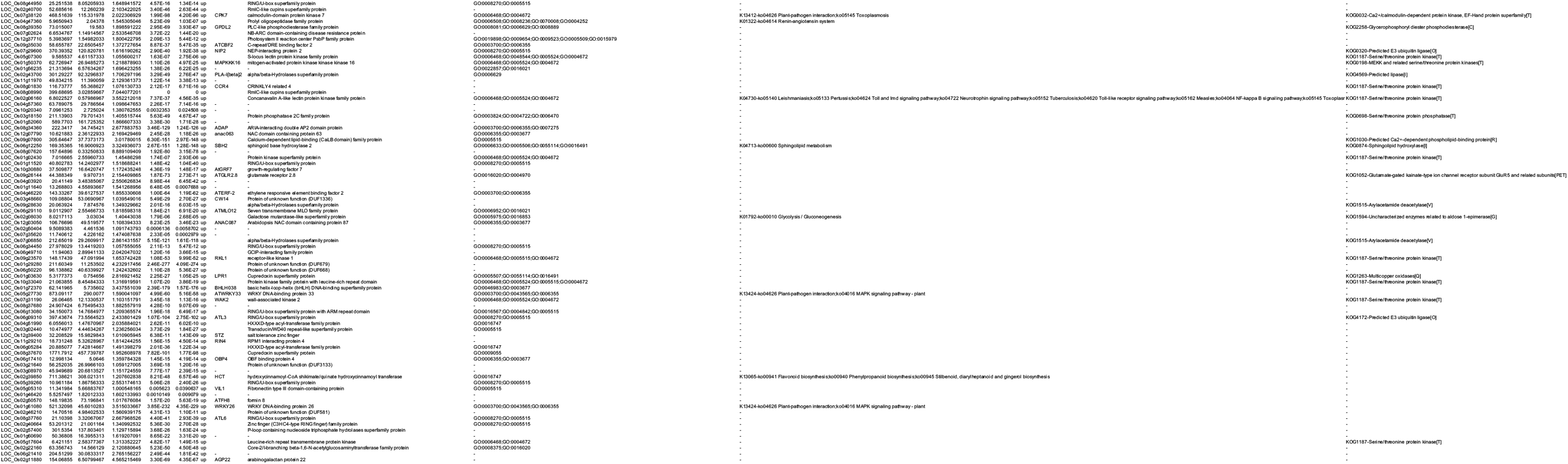
DEGs identified by chitin treatment.

**Supplimental Table 4.**
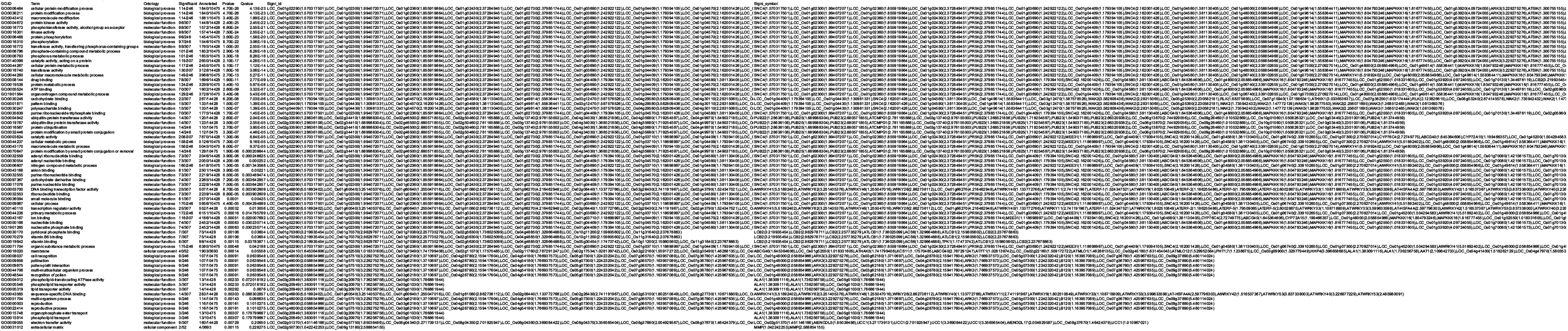
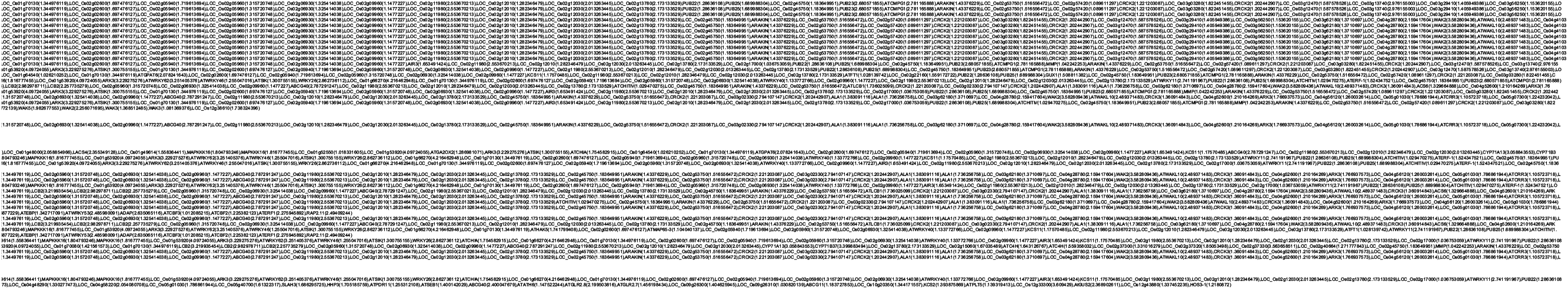
GO analysis of overlap upregulated genes in IRP and chitin treatment

**Supplimental Table 5.**
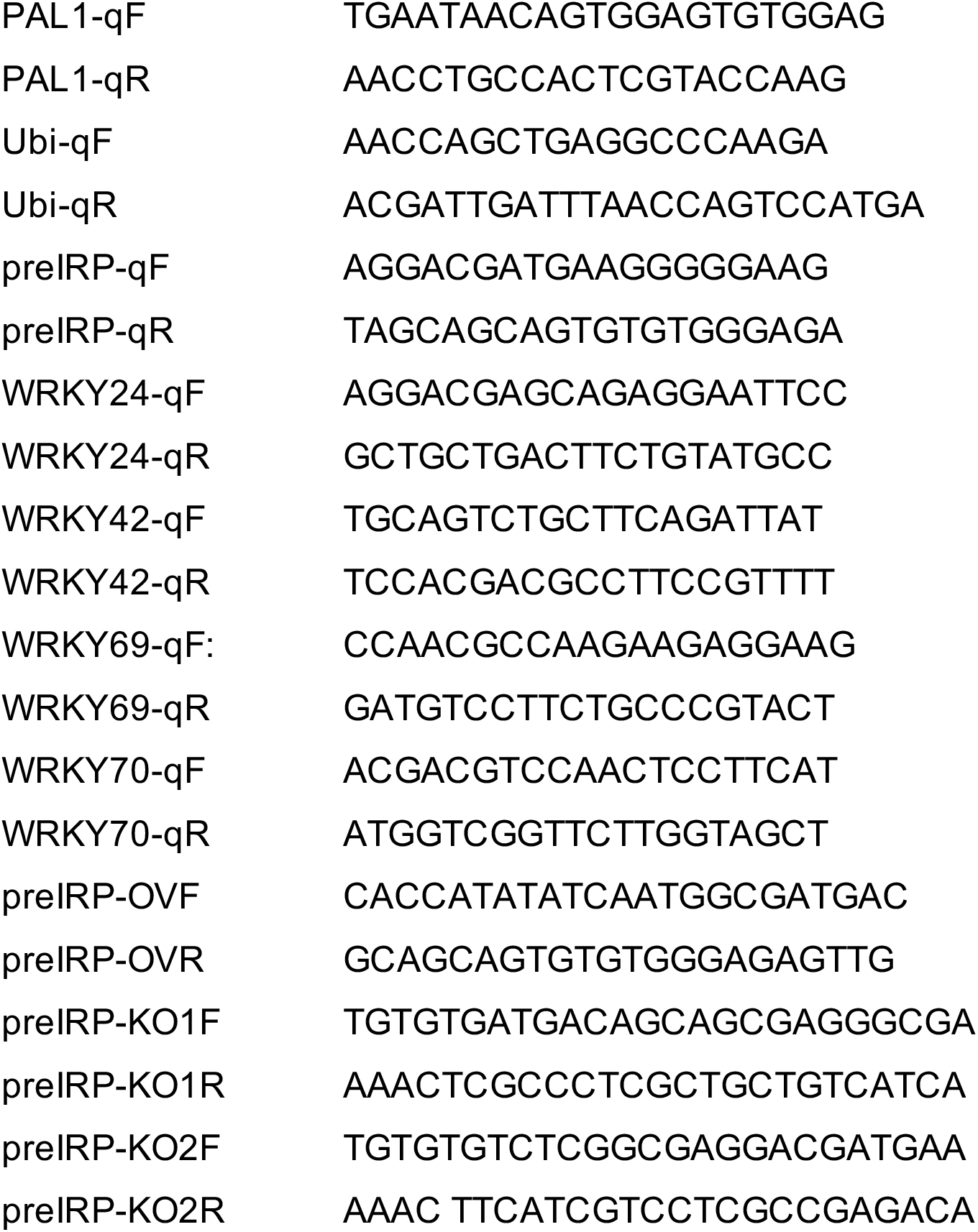
Primers for qPCR and transgenic plants.

